# Evolution of the PRD1-adenovirus lineage: a viral tree of life incongruent with the cellular universal tree of life

**DOI:** 10.1101/741942

**Authors:** Anthony C. Woo, Morgan Gaia, Julien Guglielmini, Violette Da Cunha, Patrick Forterre

## Abstract

Double-stranded DNA viruses of the PRD1-adenovirus lineage are characterized by homologous major capsid proteins containing one or two β-barrel domains known as the jelly roll folds. Most of them also share homologous packaging ATPases of the FtsK/HerA superfamily P-loop ATPases. Remarkably, members of this lineage infect hosts from the three domains of life, suggesting that viruses from this lineage could be very ancient and share a common ancestor. Here we analyzed the evolutionary history of these cosmopolitan viruses by inferring phylogenies based on single or concatenated genes. These viruses can be divided into two supergroups infecting either eukaryotes or prokaryotes. The latter can be further divided into two groups of bacterioviruses and one group of archaeoviruses. This viral tree is thus incongruent with the cellular tree of life in which Archaea are closer to Eukarya and more divergent from Bacteria. We discuss various evolutionary scenarios that could explain this paradox.

## Introduction

Studying virus origin and evolution is a challenging exercise, especially when addressing early co-evolution with cellular domains. While cellular domains (Archaea, Bacteria, and Eukarya) have been established based on ribosome sequences and more recently single-copy genes, viral lineages (or supergroups) have been suggested based on the conservation of their major capsid proteins (MCPs) structure (*1*–*4*). It has been proposed that MCP can be considered as the “viral self” or the “viral hallmark protein” (*1*, *5*, *6*). Viruses can be broadly divided into two types of lineages: domain-specific (identified in hosts from a single domain) and cosmopolitan lineages (identified in hosts from more than one domain) (*7*, *8*). To date, only viruses from the HK97 and the PRD1-adenovirus lineages, both corresponding mostly to double-stranded (ds) DNA viruses, infect host from the three domains of life (*1*, *3*, *9*, *10*). The HK97 lineage mostly consists of archaeal and bacterial viruses, whereas the viruses of the PRD1-adenovirus lineage are well represented in the virosphere associated with all three domains (archaeoviruses, bacterioviruses, and eukaryoviruses) and this lineage thus becomes an ideal subject to study the evolution of this viral lineage in the context of the universal tree of life.

The PRD1-adenovirus lineage was initially defined by the conservation of a double-jelly roll (DJR) fold in the MCP (*11*). This lineage is now characterized by either a single MCP with the DJR fold (*1*, *3*, *12*) or two MCPs, each with a single-jelly roll fold (SJR) (*13*–*16*). In both cases, the β-strands of their folds are oriented vertically relative to the capsid surface. These viruses will be called hereinafter DJR and SJR DNA viruses, respectively. The first attempt to determine the evolutionary relationships between DJR viruses was consequently based on structural alignments and confirmed that all viruses with DJR fold MCPs are indeed evolutionarily related (*11*).

The PRD1-adenovirus lineage encompasses different families as listed in Table 1. Among eukaryoviruses, all members of the PRD1-adenovirus lineage are DJR DNA viruses. Beside *Adenoviridae*, this lineage includes all members of the Nucleo-Cytoplasmic Large DNA Viruses (NCLDVs) assemblage (members of the families *Mimiviridae*, *Phycodnaviridae*, *Marseilleviridae*, *Iridoviridae*, *Ascoviridae*, *Poxviridae*, *Asfarviridae*, and related viruses), the *Lavidaviridae* (virophages related to viruses from the family of *Mimiviridae*) and the Polintoviruses (*12*, *17*). In prokaryotes, most members of the PRD1-adenovirus lineage are also DJR DNA viruses, except *Sphaerolipoviridae*, which consists of SJR DNA viruses grouping two subfamilies of archaeal viruses and one subfamily of bacterial viruses (*18*, *19*). The best-known archaeal and bacterial DJR viruses of the PRD1-adenovirus lineage are small viruses, such as *Turriviridae*, *Tectiviridae*, and *Corticoviridae*, exemplified by the Sulfolobus Turreted Icosahedral Virus (STIV), the bacteriophage PRD1, and the Pseudoalteromonas virus PM2 infecting *Sulfolobus*, *Escherichia coli*, and *Pseudoalteromonas*, respectively (*12*, *20*). Most of these prokaryotic viruses are also known to integrate into hosts genomes (*20*, *21*) or exist as free plasmids corresponding to defective viruses (*22*). In terms of representation, the prokaryotic portion of the lineage was considered scarce compared to the eukaryotic counterpart, but until recently Koonin and co-workers highlighted the unexpected diversity of these viruses in archaeal and bacterial genomes and metagenomes (*20*). The authors identified five groups of prokaryotic dsDNA viruses related to the PRD1-adenovirus lineage based on detecting sequence similarities between their MCP: Odin (named after an integrated element present in the metagenome-assembled genome of an Odinarchaeon), PM2 (with some members, the Autolykoviruses, abundant in marine microbial metagenomes(*21*)), STIV (all the viruses infecting Archaea belong to this group, with the exception of the one presumably infecting the Odinarchaeon), Bam35/Toil/Cellulomonas, and PRD1. An additional group, FLiP (*Flavobacterium*-infecting, lipid-containing phage), composed of rolling-circle ssDNA viruses with a homologous, yet very divergent MCP, has also been identified (*23*).

**Table 1.**
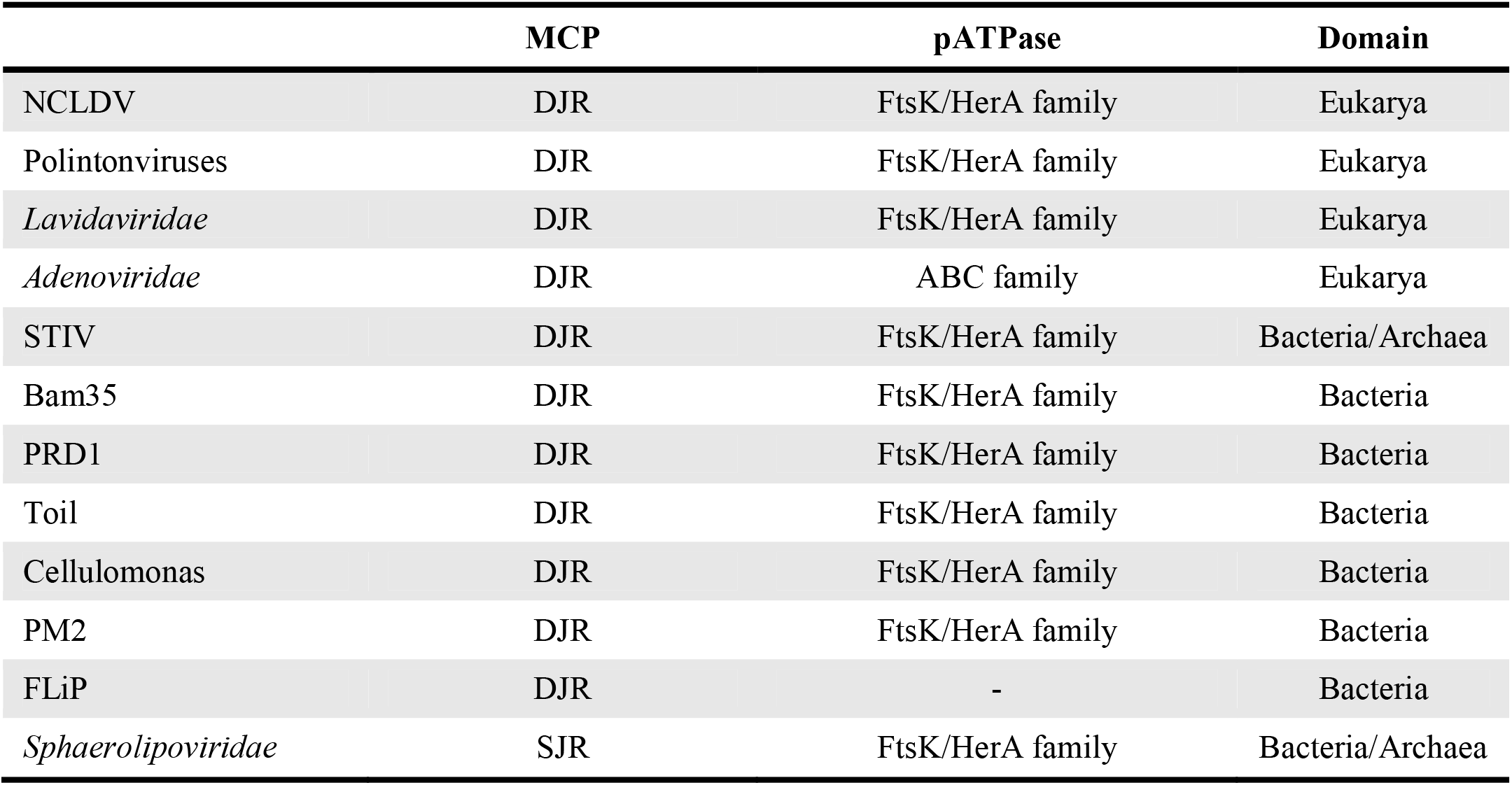
The PRD1-adenovirus lineage. This table summarizes the shared genes between different families of this lineage. Abbreviations: DJR, double jelly-roll; SJR: single jelly-roll; NCLDV, Nucleo-Cytoplasmic Large DNA Viruses; STIV, Sulfolobus turreted icosahedral virus; FLiP, Flavobacterium-infecting bacteriophage; MCP, major capsid protein; pATPase, packaging ATPase.

In a phylogenetic tree based on the common core of the MCP published in 2013, the viruses of the DJR PRD1-adenovirus were divided into three groups: one corresponding to Adenoviruses, another grouping NCLDVs (represented by two members) with *Lavidaviridae*, and the third one including viruses infecting Archaea and Bacteria (*11*). Furthermore, the evolutionary relationships among the PRD1-adenovirus lineage were recently investigated using pairwise sequence similarities between DJR viruses (*20*, *24*). These analyses demonstrated that there are evolutionary links between archaeoviruses and bacterioviruses as well as eukaryoviruses and bacterioviruses. With the advent of metagenomics approaches, many new viruses related to the PRD1-adenovirus lineage have been identified recently, providing important insights into the evolution of viruses from this lineage. This strengthens the call for a more thorough study to examine the evolutionary relationships among all MCPs from the DJR viruses.

In addition to the MCP, most members of the PRD1-adenovirus lineage, with a few exceptions, share a packaging ATPase (pATPase) of the FtsK/HerA superfamily P-loop ATPases (Table 1) (*3*, *20*). The MCP and pATPase were also part of the core genes defined in our recent study on the evolution of the NCLDV assemblage (*25*), and concatenating the two proteins proved useful in rooting the phylogenetic tree of NCLDVs with the Polintoviruses as an outgroup, suggesting that such an approach could be extended to the whole PRD1-adenovirus lineage.

Here we show that the concatenated MCP and pATPases genes of the viruses of the PRD1-adenovirus lineage produces a robust viral phylogenetic tree characterized by a major division between eukaryotic and prokaryotic viruses, in contradiction with the topology of the universal cellular tree, which is characterized by the grouping of Archaea with eukaryotes. Accordingly, the viral phylogenetic tree is not in congruence with the cellular phylogenetic trees inferred using universal markers shared between the three cellular domains. We explore various scenarios that could explain this discrepancy between the viral tree and the cellular universal tree of life.

## Results

### The MCP and the pATPase genes were used as the hallmark proteins to identify viruses from the PRD1-adenovirus lineage

We retrieved MCP and pATPase sequences using PSI-BLAST searches against the GenBank non-redundant protein sequence database (nr), including sequences recovered from proviruses (Materials and Methods). To validate the selected MCP and pATPase sequences in our dataset, we generated protein models for all selected sequences and compared these predicted structures to the PDB database. The DJR MCPs associated to groups and families previously described indeed matched their corresponding structures in the public databases, except for the putative MCPs identified from the Odin group, confirming the structural conservation of the motif in the MCP of most of the viruses from the PRD1-adenovirus lineage (fig. S1 and data file S1). The results also showed that MCPs from the SJR groups were very divergent from those of the DJR groups. The MCP from *Adenoviridae* was unique in exhibiting several additional structural elements (fig. S1). Our analysis also showed that all pATPases share similar predicted structures including this time those from the SJR groups. The only exception was the pATPase of the *Adenoviridae*, which had again, like in the MCP analysis, the most distant structure compared to the other groups of DJR viruses (fig. S2 and data file S1).

A recent study has used the MCP genes of the PRD1-adenovirus lineage to create amino acid sequence similarity networks to analyze evolutionary relationships between viruses from this lineage (*20*). We thus performed a similar analysis based on the pATPases (fig. S3). Almost all viruses of the PRD1-adenovirus lineage were clustered together when using the pATPases for the analysis, except for the *Adenoviridae*. This corroborates previous observations based on amino-acid signatures and secondary structure prediction suggesting that pATPases of the *Adenoviridae* were not specifically related to other pATPases of viruses of the PRD1-adenovirus lineage but were derived from ATPases of the ABC superfamily (*26*), indicating an exchange of pATPases during the evolution of this viral family. Surprisingly, we found that the pATPases of the SJR DNA viruses did not form a distinct cluster, as expected for an outgroup, but strongly clustered together with the pATPases of DJR archaeoviruses and bacterioviruses. It was possible to distinguish two large assemblages in the pATPase network, one corresponding to a core NCLDVs loosely associated to the *Asfarviridae* on one side and the *Poxviridae* and the Polintoviruses on the other, and a second one corresponding to archaeoviruses and bacterioviruses (including SJR viruses), loosely associated to the *Lavidaviridae*.

### Phylogenetic positions of the PRD1-adenovirus lineage viruses inferred from the single protein genes

The result of the sequence similarity network suggested that the pATPase gene could be a good marker for delineating the phylogeny of the viruses from the PRD1-adenovirus lineage, in addition to the MCP that was used in recent studies (*11*, *20*). Sequences of the MCPs (excluding and both SJR viruses and the *Adenoviridae*) and pATPases were aligned and trimmed (Materials and Methods). Phylogenetic analyses were performed separately on the MCP and the pATPase using the resulting datasets within the maximum likelihood (ML) framework. DJR eukaryoviruses were clearly separated from archaeoviruses and bacterioviruses in both trees, in agreement with previous observations made using pairwise sequence similarities or comparative structural analyses (*11*, *24*). In particular, unlike the result obtained with the sequence similarity network, members of the *Lavidaviridae* were grouped with other eukaryoviruses in the pATPase tree. We thus decided to root the trees between “prokaryotic” and “eukaryotic” DJR viruses. As a result, DJR eukaryoviruses and viruses infecting Archaea or Bacteria formed two monophyletic clades. Moreover, we recovered the monophyly of most previously defined assemblages at various taxonomic ranks (groups, families or superfamilies). The few exceptions were the STIV, the *Phycodnaviridae* and the *Iridoviridae* that were paraphyletic in both the MCP and pATPase trees, and the PM2 group which was paraphyletic in the pATPase tree (fig. S4 and S5). Mollivirus, which is related to pandoraviruses, branched within the *Phycodnaviridae* and this corroborates previous observations (*25*, *27*).

In the MCP tree, the NCLDV formed a monophyletic sister group to the *Lavidaviridae*, whereas in the pATPase tree, the *Lavidaviridae* branched at the base of the eukaryotic clade and the NCLDV were not monophyletic as the *Poxviridae* were sister group to the Polintoviruses. On the prokaryotic side, viruses of the STIV group were the sister group to viruses of the PM2 group (except for the two *Sulfolobus* STIV), whereas the five other groups of DJR bacterioviruses (the Bam35, the PRD1, the Toil, the Cellulomonas and the FLiP) emerged at the base of this part of the tree (fig. S4).

In the pATPase tree, the Bam35, the PRD1, the Toil and the Cellulomonas groups formed a monophyletic cluster that was sister group to all STIV (except two members of the STIV group) whereas the PM2 group and the two STIV members branched at the base of the tree, within SJR viruses of the *Sphaerolipoviridae* family (fig. S5). Remarkably, the SJR pATPases did not form a monophyletic group well separated from DJR viruses, as expected if DJR viruses originated from SJR viruses (*28*), but grouped with DJR viruses infecting prokaryotes in agreement with the network analysis (fig. S3 and S5). To better address the evolutionary relationship between the pATPases of SJR and DJR viruses, we performed another phylogenetic analysis using only sequences of bacterial and archaeal viruses (fig. S6). Interestingly, the pATPases of archaeal SJR DNA viruses branched within the clade of archaeal DJR DNA viruses, whereas the pATPases of bacterial SJR viruses formed a separate clade. It has been previously suggested that DJR viruses originated from the simpler SJR type by ancestral gene duplication of one of the two SJR fold MCPs that possibly occurred before the emergence of the Last Universal Common Ancestor (LUCA) (*29*, *30*). If one assumes that SJR viruses, defined by their very divergent MCPs, predated DJR viruses as previously proposed (*28*), this suggests that they replaced later on their pATPases with those of DJR viruses infecting the same hosts. Alternatively, SJR viruses might have emerged twice independently from DJR viruses and the MCP genes of SJR viruses might have diverged rapidly following their structural rearrangements.

To better compare the MCP and pATPase trees, we removed the sequences of the SJR viruses whose MCP cannot be included in the MCP tree from the pATPase dataset and we removed viruses of the FLiP group that have no detectable pATPases from the MCP dataset (Fig. 1 and fig. S7 and S8). The removal of SJR did not change the topology of the pATPase tree, except for some modifications in the relative branching of DJR viruses from the PM2 and the STIV groups. Surprisingly, the removal of the FLiP viruses from the MCP tree also introduced some modifications in the relationships between the different groups of viruses infecting eukaryotes, especially the *Poxviridae* and the *Asfarviridae*. The *Poxviridae* and the *Asfarviridae* are however known to be unstable taxa, possibly due to their frequent long branches. Nevertheless, most of the clades were still monophyletic and the bipartition corresponding to eukaryoviruses and prokaryoviruses was still present.

**Fig. 1.**
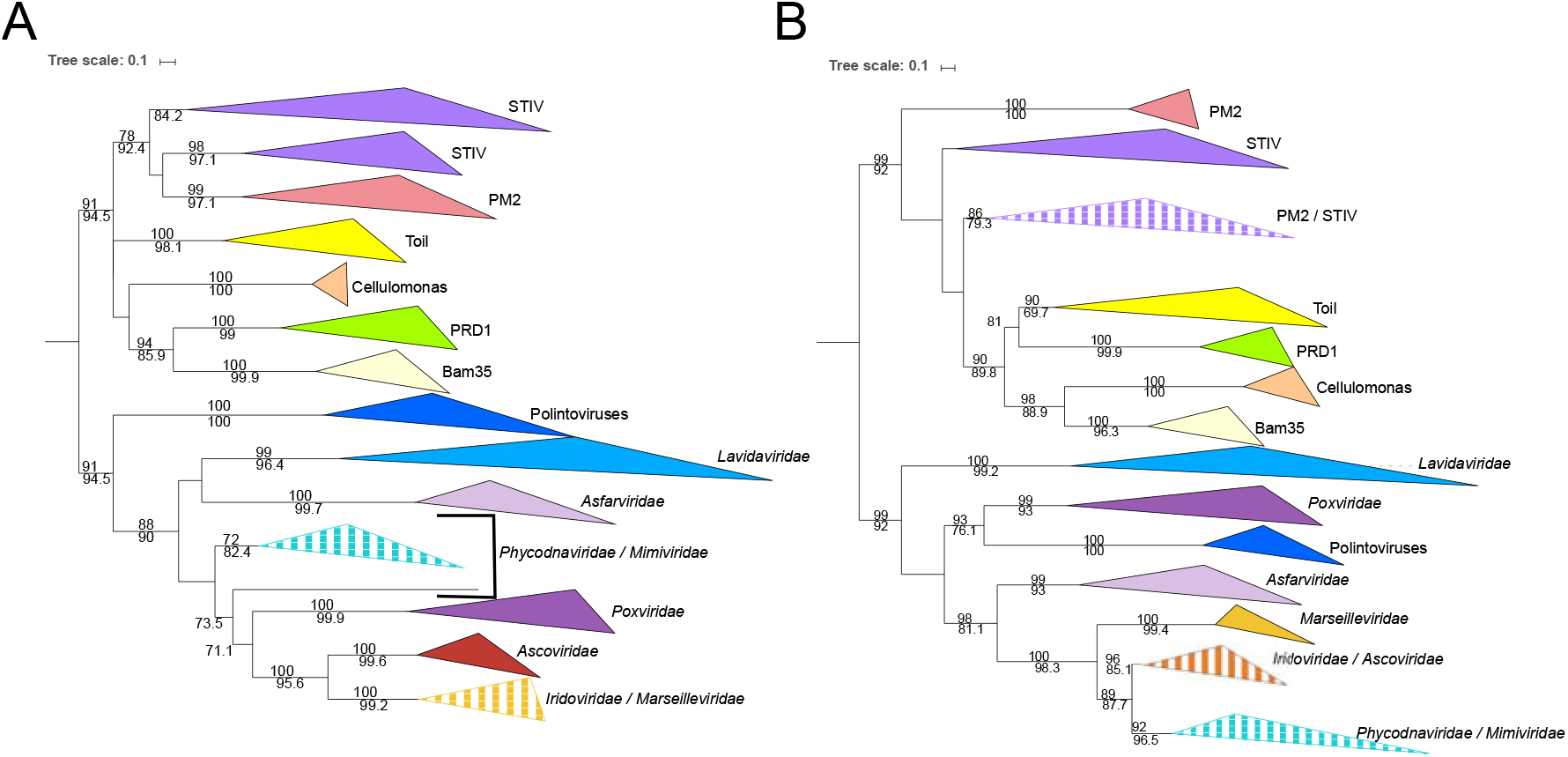
Maximum likelihood (ML) single-protein trees of the two hallmark proteins of the viruses from the PRD1-adenovirus lineage. ML phylogenetic trees of (**A**) the major capsid protein (MCP) and (**B**) packaging ATPase (pATPase). The root of the ML phylogenetic tree was between the prokaryotic and eukaryotic members. The scale-bar indicates the average number of substitutions per site. Values on top and below branches represent support calculated by ultrafast bootstrap approximation (UFBoot; 1,000 replicates) and SH-like approximate likelihood ratio test (aLRT; 1,000 replicates), respectively. Only values superior to 70 are shown. The best-fit model for the MCP tree was LG + F + R4, which was chosen according to Bayesian Information Criterion (BIC) and the alignment has 103 sequences with 237 positions. The best-fit model for the pATPase tree was LG + R6, which was chosen according to Bayesian Information Criterion (BIC) and he alignment has 103 sequences with 171 positions. More detailed versions of the trees are shown in fig. S7 and S8.

### Concatenation of the MCP and the pATPase genes highlights an unexpected evolutionary scenario

The small differences observed between the MCP and the pATPase trees, the uncertainties in the topologies due to the removal of some sequences, and the low support values of most branches in these single trees were anticipated considering the divergence of the analyzed proteins. However, these trees also exhibited some clear-cut congruence, such as the eukaryotic/prokaryotic split, the monophyly of the NCLDVs other than the *Poxviridae*, the monophyly of several previously defined groups of viruses infecting Archaea and Bacteria, the grouping of the STIV and the PM2 on one side and the grouping of the PRD1, the Bam35, the PRD1, the Toil, and the Cellulomonas on the other (Fig.1 and fig. S7 and S8). This suggested that we could concatenate the MCP and the pATPase sequences to obtain a more reliable phylogeny. We adopted the new version of transfer bootstrap in which the presence of inferred branches in replications is measured using a gradual ‘transfer’ distance and the resulting supports do not induce falsely supported branches (*31*). In the concatenated ML tree, we recovered the division between DJR viruses infecting eukaryotes and those infecting prokaryotes, as well as the monophyly of all previously defined groups of DJR viruses, except for the STIV group. Interestingly, the STIV group was now divided into two subgroups corresponding to DJR viruses infecting either Archaea or Bacteria and most of the branch supports were above 0.9 (in the range of 0 to 1), indicating that the tree is very robust (Fig. 2).

**Fig. 2.**
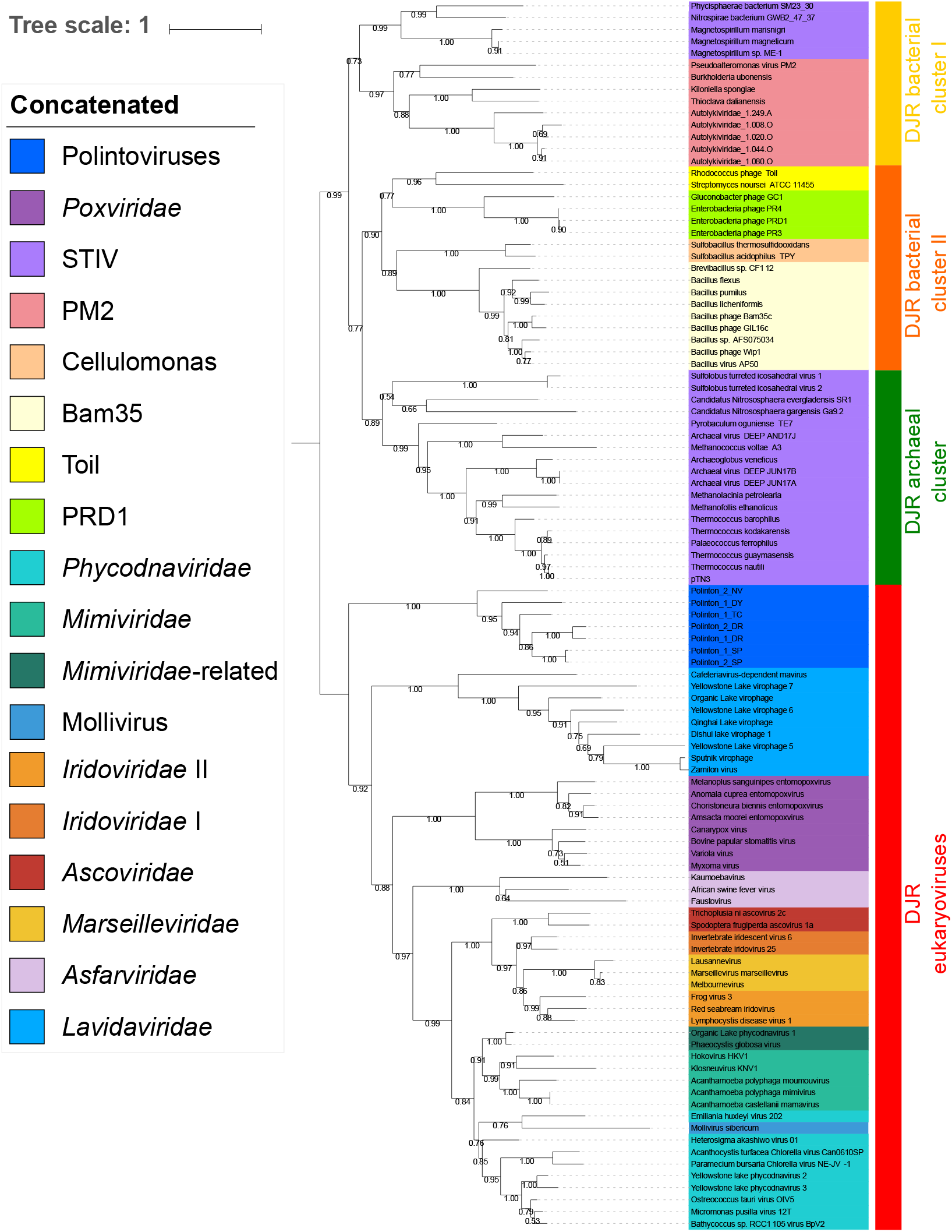
Maximum likelihood (ML) phylogenetic tree of the concatenated MCP and pATPase genes. The ML phylogenetic tree was rooted between the prokaryotic and eukaryotic members. The scale-bar indicates the average number of substitutions per site. Values on the branches represent transfer bootstrap expectation (TBE) support calculated by nonparametric bootstrap (1,000 replicates). The best-fit model was LG + F + R6, which was chosen according to Bayesian Information Criterion (BIC). The alignment has 107 sequences with 408 columns.

Among the eukaryoviruses of the tree, the Polintoviruses were basal to all eukaryotic groups whereas the NCLDV assemblage was sister group to the *Lavidaviridae*. The *Poxviridae* were the first NCLDVs branching family, followed by the *Asfarviridae*. Notably, we recovered the two major groups of NCLDVs that we previously identified based on 8 marker core genes (*25*), the MAPI (*Marseilleviridae*, *Ascoviridae*, Pitho-like viruses, and *Iridoviridae*) and the PAM (*Phycodnaviridae, Asfarviridae* and the recently proposed order “Megavirales”), except that the *Asfarviridae* were sister group to these two superclades instead to be part of the PAM and the *Phycodnaviridae* were paraphyletic. Remarkably, we also previously obtained *Asfarviridae* as sister group to other NCLDVs in a tree based on the two large RNA polymerase subunits, using cellular sequences as an outgroup (*25*). We previously noticed that the *Poxviridae* had exceptionally long branches and variable positions in single-gene trees of NCLDV proteins. In particular, they tended to attract the long branches of the *Asfarviridae* in our previous analyses. We thus removed both the *Poxviridae* and the *Asfarviridae* from the dataset and we obtained a tree in which NCLDVs are no more sister group to the *Lavidaviridae* (fig. S9), but the Polintoviruses, as previously proposed based on phenotypic analyses (*32*).

Noticeably, two strongly supported clades of viruses infecting bacteria and one clade of viruses infecting Archaea were recovered on the prokaryotic side of the tree (Fig. 2). The bacterial members of the STIV formed a strongly supported monophyletic clade with bacterial viruses/proviruses of the PM2 group that branched at the base of this clade and will be hereinafter called the DJR bacterial cluster I. The second bacterial clade with the Cellulomonas, the Bam, the Toil and the PRD1 (hereinafter called the DJR bacterial cluster II) was similar to a clade first observed with the pATPase tree. The third clade included all archaeal members of the STIV group (hereinafter called the DJR archaeal cluster). The DJR archaeal cluster became sister group to the DJR bacterial cluster I when both the *Poxviridae* and the *Asfarvirida*e were removed from the analysis, with members the DJR bacterial cluster II becoming paraphyletic at the base of the “prokaryotic clade” (fig. S9).

The monophyly of DJR archaeoviruses, previously observed with the complete MCP tree was only weakly supported in the concatenated tree with or without the *Poxviridae* and the *Asfarvirida*e. To improve the resolution, we removed eukaryoviruses from the concatenated tree. We again obtained the monophyly of the DJR bacterial clusters I, II, and the archaeal cluster if we rooted the tree between Archaea and Bacteria (Fig. 3). On this occasion, we obtained good support for the monophyly of Archaea. Interestingly, viruses infecting Euryarchaea and those infecting Crenarchaea or Thaumarchaea were clearly separated in the tree, suggesting that these viruses might have been present at the time of the last common ancestor of Archaea and later on co-evolved with cellular hosts from this domain.

**Fig. 3.**
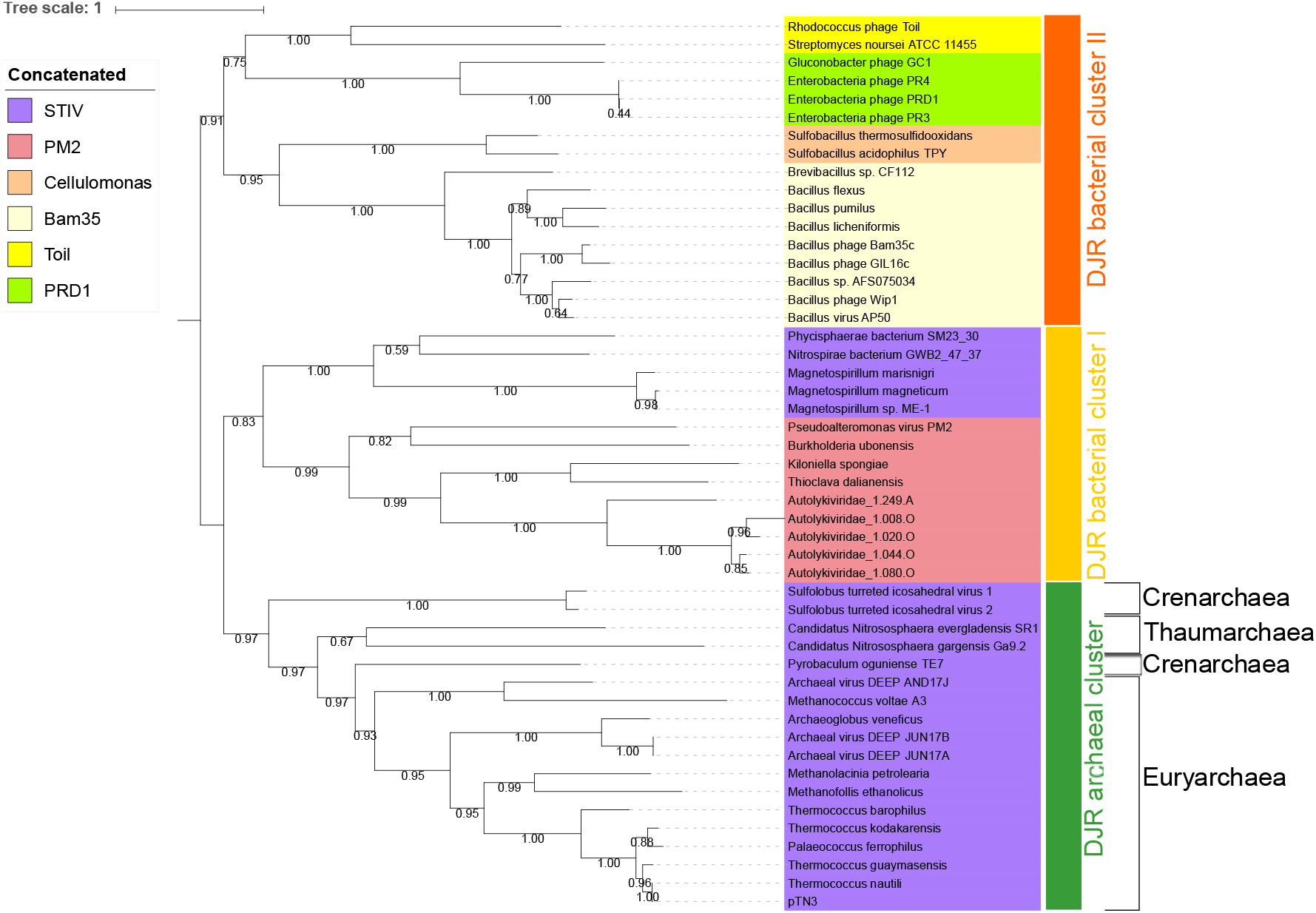
Maximum likelihood (ML) phylogenetic tree of the concatenated MCP and pATPase genes of bacterioviruses and archaeoviruses. The ML phylogenetic tree was rooted with the DJR bacterial cluster II. The scale-bar indicates the average number of substitutions per site. Values on the branches represent transfer bootstrap expectation (TBE) support calculated by nonparametric bootstrap (1,000 replicates). The best-fit model was LG + R4, which was chosen according to Bayesian Information Criterion (BIC). The alignment has 49 sequences with 468 columns.

## Discussion

The PRD1-adenovirus lineage is one of the two major viral lineages with members infecting hosts from all three domains of life. It has long been thought that inferring phylogeny of this lineage was out of reach because of the absence of significant sequence similarities between their MCPs (*11*). This could be because the first viruses identified in this lineage, PRD1 and Adenoviruses, are the most divergent viruses of this lineage. However, many additional large groups of DNA viruses from the three domains of life were later on progressively incorporated in this lineage, revealing previously undetectable connection at the primary sequence level between their MCPs and pATPases. Here, we show that most members of this lineage can be placed in a robust “universal tree” based on the concatenation of these two proteins. A similar strategy has been recently adopted to produce a global evolutionary history of RNA viruses based on the phylogeny of their RNA-dependent RNA polymerases (*33*). Our results are supported by the fact that we recovered most groups previously defined on different criteria, with the only exception of the STIV group (*20*). In particular, the internal phylogeny of the NCLDVs is comparable to the one that we previously obtained with eight-core genes (*25*).

The tree presented here will certainly change in the future with the discovery of new viral groups belonging to this lineage and the discovery of new members of the existing groups. Nevertheless, our findings suggest that the phylogenetic tree of the concatenated MCP and pATPase can be used as a backbone to review current hypotheses about the evolution of this lineage and propose new ones.

Interestingly, our in-depth phylogenetic analyses reveal a great divide between DJR viruses infecting prokaryotes (Archaea and Bacteria) and those infecting eukaryotes. This produces a “viral tree of life” strikingly different from the cellular tree based on universal proteins in which Archaea and eukaryotes are closely related (*34*–*37*). Interestingly, the topology of this viral tree of life corroborates phenotypic observations showing that the mobilomes of Archaea and Bacteria are both very similar and strikingly different from the eukaryotic mobilome (*38*). For instance, Caudovirales (head and tailed “phages”), conjugative plasmids, casposons, or several families of IS elements are only present in Archaea and Bacteria (*39*–*44*), whereas several families of RNA viruses, retroviruses, and certain families of IS elements are specific to eukaryotes (*38*, *45*). Similarly, defense systems are also distributed according to the “eukaryote/prokaryote” divide, with the CRISPR-Cas system or the toxin-antitoxin systems only present in Archaea and Bacteria whereas the RNAi system only exists in eukaryotes (*46*, *47*).

It has been suggested that all eukaryoviruses, including those from the PRD1-adenovirus lineage, originated from a melting pot of bacteriophages (bacterioviruses) infecting the bacterium at the origin of mitochondria (*48*). Regarding the origin of DJR viruses, it was suggested that Polintoviruses evolved from a *Tectiviridae* and the Polintoviruses subsequently became the ancestor of *Lavidaviridae* and NCLDVs (*32*). However, in that scenario, DJR eukaryoviruses, especially Polintoviruses, should have been more closely related to *Tectiviridae* than to other DJR bacterioviruses in the MCP/pATPase DJR tree, which is not the case.

Phylogenetic analyses of cellular and NCLDV RNA polymerases revealed that NCLDVs had already diverged before the Last Eukaryotic Common Ancestor (LECA) (*25*). Our present study indicates that the divergence between NCLDVs, Polintoviruses and *Lavidavirida*e took place even earlier. This renders unlikely that the transformation of an ancestral prokaryotic-like DJR virus into these three groups of eukaryoviruses had enough time to occur during the evolutionary period between the first eukaryote with a proto-mitochondrion and LECA. If they all originated from the mitochondrion, it would leave a rather short lapse of time before LECA for so many various families to emerge. Notably, among these three groups of eukaryoviruses, one group encompasses diverse large and giant viruses infecting a range of host from unicellular algae to human beings. A more plausible related hypothesis could be that DJR eukaryoviruses evolved from DJR viruses that infected prokaryotes that entered into symbiosis with proto-eukaryotes much before the final endosymbiosis that led to mitochondria, leaving more time for the evolution and diversification of DJR eukaryoviruses.

If DJR eukaryoviruses originated indeed from bacterioviruses or archaeoviruses (hereinafter called the “prokaryotic” ancestor hypothesis), the DJR universal tree could be rooted with the DJR archaeal cluster (fig. S10), the DJR bacterial clusters (fig. S11 and S12) or between the DJR archaeal cluster and the DJR bacterial cluster I (Fig. 4A and fig. S13). In this scenario, according to our MCP/pATPase tree, DJR eukaryoviruses could have originated from a third unknown group of DJR viruses, sister group to DJR bacterial cluster I. In any case, it remains unclear in the “prokaryotic” ancestor hypothesis how a virus infecting a prokaryotic symbiont could have acquired the capacity to infect the proto-eukaryotic host.

**Fig. 4.**
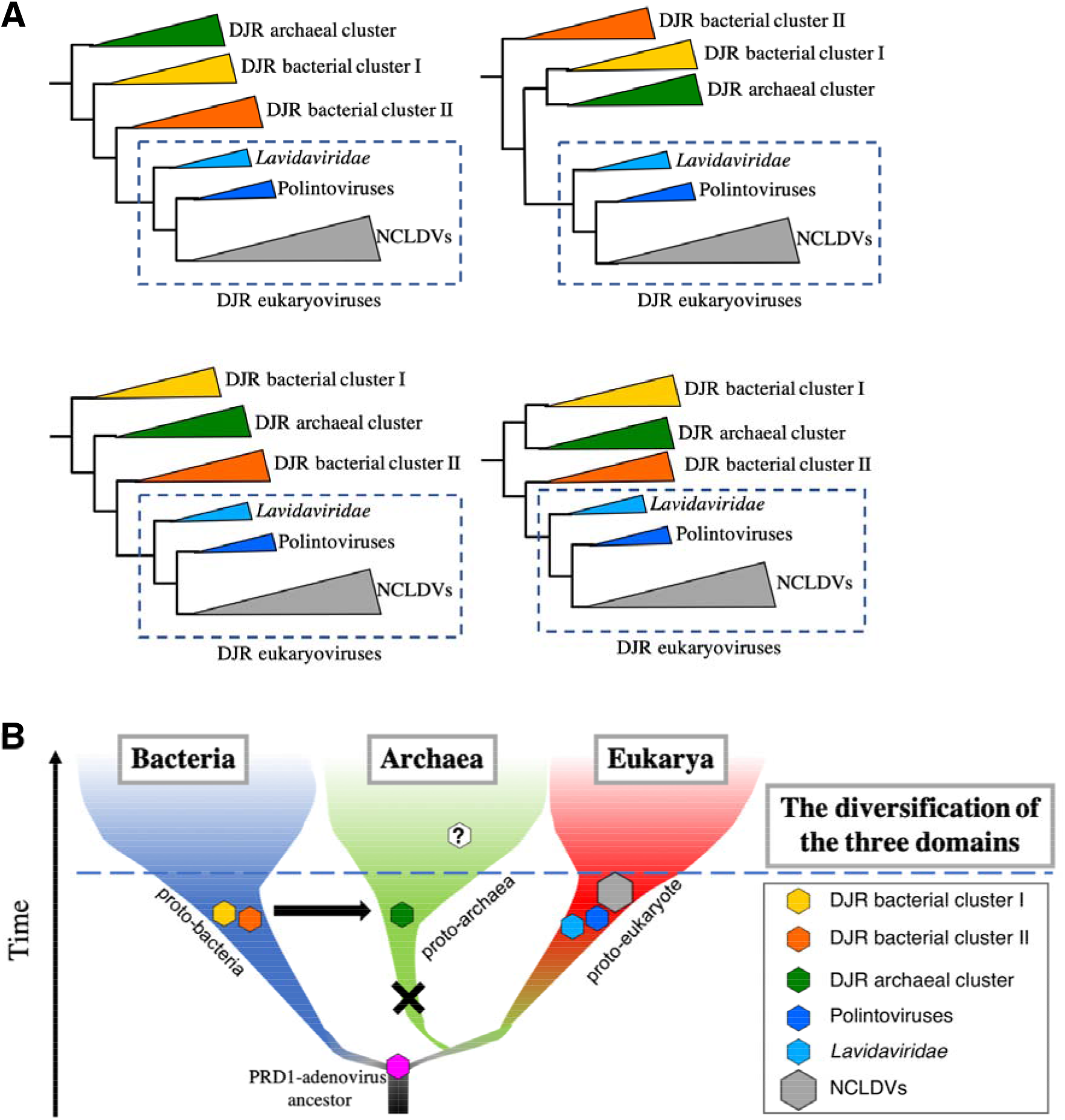
Hypothetical scenarios for the evolution of viruses of the PRD1-adenovirus lineage. (**A**) “Prokaryotic” ancestor hypothesis. Depending on the rooting, different possible topologies of the double jelly-roll viruses of this lineage are depicted (see fig. S10, S11, S12 and S13). In any case, double jelly-roll (DJR) eukaryoviruses originated from viruses, which were closely related to DJR bacterial cluster. (**B**) The enigma of the PRD1-adenovirus lineage. The presence of viruses from the PRD1-adenovirus lineage across the three domains of life suggests that these viruses could be very ancient. In this scenario, modern DJR archaeoviruses are more closely related to DJR bacterioviruses. Ancient viruses which could infect archaea were lost after the divergence of Archaea and Eukarya. After that, modern DJR archaeoviruses were being transferred from DJR bacterioviruses before the diversification of Archaea. Another possibility is some of the viruses which are closely related to DJR eukaryoviruses have yet to be found.

The presence of DJR viruses in the three domains has been often used as an argument suggesting that these viruses were already present at the time of the Last Universal Common Ancestor (LUCA) (*1*)(*49*)(*50*). In the “prokaryotic” ancestor hypothesis, the presence of DJR viruses in LUCA favors rooting the whole DJR tree between archaeoviruses and bacterioviruses, as in fig. S13. However, the branch between archaeoviruses and bacterioviruses was very short in fig. S13, whereas the branch between archaeal and bacterial proteins is always very long in the trees of universal proteins that were present in LUCA (*34*)(*51*). In the “prokaryotic” ancestor hypothesis, it thus seems more likely that DJR viruses originated after LUCA, either in proto-archaea or proto-bacteria and were later on transferred from one domain to the other.

Alternatively, the presence of DJR viruses in LUCA would explain the clear-cut separation of prokaryotic and eukaryotic DJR viruses in the whole DJR tree. However, considering that viruses generally co-evolve with their hosts, it remains obscure why DJR archaeoviruses are much more similar to DJR bacterioviruses than to DJR eukaryoviruses, since Archaea and Eukarya are closely related in the universal tree of life. A plausible explanation for this paradox could be that eukaryotic-like DJR viruses were first lost in the stem lineage of Archaea and reintroduced later on in this domain from Bacteria (or proto-bacteria) before the diversification of Archaea (Fig. 4B). The present absence of eukaryotic-like DJR viruses in Archaea could be also due to a sampling bias, some of these archaeal viruses being still hidden in some unexplored places around the globe.

The presence of DJR viruses in the three cellular domains thus raises challenging questions about their origin and evolution. Here, we have shown that phylogenies based on the concatenation of their MCP and pATPase not only can help to formulate more precisely previous evolutionary hypotheses but also to propose new ones, such as the monophyly of DJR archaeoviruses and to define new assemblages, such as DJR clusters I and II in Bacteria. Future identification and isolation of new viral families in the PRD1-adenovirus lineage will be essential to provide insights into various scenarios for the history of this lineage.

## Materials and Methods

### Selection of MCP/pATPase sequences

Representative MCP/pATPase sequences from different groups of the DJR viruses were used as queries for PSI-BLAST (*52*) searches against the GenBank non-redundant protein sequence database (nr). The query sequences are listed below:

| Group | Name | MCP | pATPase |
| --- | --- | --- | --- |
| *Lavidaviridae* | Sputnik virophage | YP_002122381 | YP_002122364 |
| *Adenoviridae* | Frog adenovirus 1 | NP_062443 | NP_062434 |
| STIV | Sulfolobus turreted icosahedral virus 1 | YP_025022 | YP_025021 |
| Bam35 | Bacillus phage Bam35c | NP_943764 | NP_943760 |
| PRD1 | Enterobacteria phage PRD1 | NP_040692 | NP_040689 |
| Toil | Rhodococcus phage Toil | ARK07697 | ARK07695 |
| PM2 | Pseudoalteromonas phage PM2 | NP_049903 | NP_049900 |
| FLiP | Flavobacterium phage FLiP | ASQ41214 | - |

The MCP/pATPase sequences of the NCLDVs were retrieved from a previous study that we conducted (*25*). The Polintoviruses sequences were gathered from the Repbase collection (*53*) (http://www.girinst.org/Repbase_Update.html): Polinton-2_NV, Polinton-1_DY, Polinton-1_TC, Polinton-1_SP, Polinton-2_SP, Polinton-2_DR, Polinton-1_DR. Finally, the sequences from the SJR group were recovered based on previously identified sequences (*19*).

Putative MCP/pATPase sequences were aligned with the query sequences for the examination of the conserved structural elements using MAFFT (*54*). Prediction of the secondary structure was performed using Phyre2 (*55*) and the predicted protein structures were visualized using UCSF Chimera (*56*). The sequences used in this study are shown in the supplementary file. After removing sequences with no significant matches or low confidence levels, we obtained two different datasets of 145 and 128 sequences for the MCP and pATPase respectively.

### Network analysis

After performing the structural protein prediction analysis, all-against-all blastp analyses were performed on the refined MCP and pATPase datasets. The all-against-all integrases BlastP results were grouped using the SiLiX (for *SIngle LInkage Clustering of Sequences*) package v1.2.8 (http://lbbe.univ-lyon1.fr/SiLiX)(*57*). This approach for the clustering of homologous sequences is based on single transitive links with alignment coverage constraints. Several different criteria can be used separately or in combination to infer homology (percentage of identity, alignment score or E-value, alignment coverage). The MCP and pATPase sequences were clustered independently by similarity using SiLiX with the expect threshold of 0.001 as previously used for MCP analysis (*20*). The clustering results were analyzed and visualized using the igraph package of the R programming language (https://igraph.org/).

### Phylogenetic analysis

The alignments of the MCP sequences were performed using MAFFT v7.392 with the E-INS-i algorithm (*54*), which can align sequences with several conserved motifs embedded in long unalignable regions, whereas pATPase sequences were aligned using MAFFT with the L-INS-i algorithm (*54*), which can align a set of sequences containing sequences flanking around one alignable domain. Positions containing more than 30% of gaps were trimmed using goalign v0.2.8 (https://github.com/evolbioinfo/goalign).

### Maximum likelihood phylogenies

Single protein and concatenated protein phylogenies were conducted within the Maximum Likelihood (ML) framework using IQ-TREE v1.6.3 (*58*). We first performed a model test with the Bayesian Information Criterion (BIC) by including protein mixture models (*59*). For mixture model analyses, we used the PMSF models (*60*). The support values were either transfer bootstrap expectation (TBE) support (*31*) computed from 1000 bootstrap replicates in the case of nonparametric bootstrap using gotree v0.3.0 (https://github.com/evolbioinfo/gotree), or from 1,000 replicates for SH-like approximation likelihood ratio test (aLRT) (*61*) and ultrafast bootstrap approximation (UFBoot) (*62*).

### Visualization

The phylogenetic trees were visualized with FigTree v1.4.3 (http://tree.bio.ed.ac.uk/software/figtree/) and iTOL (*63*).

## Supplementary Materials

Fig. S1. Major capsid proteins of the PRD1-adenovirus lineage.

Fig. S2. Packaging ATPases of the PRD1-adenovirus lineage.

Fig. S3. Sequence similarity networks of the PRD1-adenovirus packaging ATPase proteins.

Fig. S4. Maximum likelihood (ML) phylogenetic tree of the major capsid protein gene of the viruses from the PRD1-adenovirus lineage.

Fig. S5. Maximum likelihood (ML) phylogenetic tree of the packaging ATPase gene of the viruses from the PRD1-adenovirus lineage.

Fig. S6. Maximum likelihood (ML) phylogenetic tree of the packaging ATPase gene of the archaeoviruses and bacterioviruses from the PRD1-adenovirus lineage.

Fig. S7. Maximum likelihood (ML) phylogenetic tree of the packaging ATPase gene of the double jelly-roll viruses from the PRD1-adenovirus lineage, excluding single jelly-roll viruses.

Fig. S8. Maximum likelihood (ML) phylogenetic tree of the major capsid protein gene of the double jelly-roll viruses from the PRD1-adenovirus lineage, excluding the FLiP members.

Fig. S9. Maximum likelihood (ML) phylogenetic tree of the concatenated major capsid protein and packaging ATPase genes, excluding the *Poxviridae* and the *Asfarviridae*.

Fig. S10. Maximum likelihood phylogenetic tree of the concatenated major capsid protein and packaging ATPase genes, rooting with the DJR archaeal cluster.

Fig. S11. Maximum likelihood phylogenetic tree of the concatenated major capsid protein and packaging ATPase genes, rooting with the DJR bacterial cluster II.

Fig. S12. Maximum likelihood phylogenetic tree of the concatenated major capsid protein and packaging ATPase genes, rooting with the DJR cluster I.

Fig. S13. Maximum likelihood phylogenetic tree of the concatenated major capsid protein and packaging ATPase genes, rooting between the DJR archaeal cluster and the DJR bacterial cluster II.

Data file S1. Results of the predicted structures of the major capsid protein and the packaging ATPase genes.

## General

This work used the computational and storage services (TARS cluster) provided by the IT department at Institut Pasteur, Paris.

## Author contributions

A.C.W. and P.F. designed the study. A.C.W., J.G., and V.D.C. performed bioinformatics experiments. A.C.W., J.G., M.G., V.D.C., and P.F. analyzed and interpreted the results. A.C.W., M.G., V.D.C. and P.F. wrote the manuscript.

## Data and materials availability

All data needed to evaluate the conclusions in the paper are present in the paper and/or the Supplementary Materials. Additional data related to this paper may be requested from the authors.

