## Supplementary file for "Evolution of the PRD1-adenovirus lineage: a viral tree of life incongruent with the cellular universal tree of life"

**This pdf file includes:**

Fig. S1. Major capsid proteins of the PRD1-adenovirus lineage.
Fig. S2. Packaging ATPases of the PRD1-adenovirus lineage.
Fig. S3. Sequence similarity networks of the PRD1-adenovirus packaging ATPase proteins.
Fig. S4. Maximum likelihood (ML) phylogenetic tree of the major capsid protein gene of the viruses from the PRD1-adenovirus lineage.
Fig. S5. Maximum likelihood (ML) phylogenetic tree of the packaging ATPase gene of the viruses from the PRD1-adenovirus lineage.
Fig. S6. Maximum likelihood (ML) phylogenetic tree of the packaging ATPase gene of the archaeoviruses and bacterioviruses from the PRD1-adenovirus lineage.
Fig. S7. Maximum likelihood (ML) phylogenetic tree of the packaging ATPase gene of the double jelly-roll viruses from the PRD1-adenovirus lineage, excluding single jelly-roll viruses.
Fig. S8. Maximum likelihood (ML) phylogenetic tree of the major capsid protein gene of the double jelly-roll viruses from the PRD1-adenovirus lineage, excluding the FLiP members.
Fig. S9. Maximum likelihood (ML) phylogenetic tree of the concatenated major capsid protein and packaging ATPase genes, excluding the *Poxviridae* and the *Asfarviridae*.
Fig. S10. Maximum likelihood phylogenetic tree of the concatenated major capsid protein and packaging ATPase genes, rooting with the DJR archaeal cluster.
Fig. S11. Maximum likelihood phylogenetic tree of the concatenated major capsid protein and packaging ATPase genes, rooting with the DJR bacterial cluster II.
Fig. S12. Maximum likelihood phylogenetic tree of the concatenated major capsid protein and packaging ATPase genes, rooting with the DJR cluster I.
Fig. S13. Maximum likelihood phylogenetic tree of the concatenated major capsid protein and packaging ATPase genes, rooting between the DJR archaeal cluster and the DJR bacterial cluster II.
Data file S1. Results of the predicted structures of the major capsid protein and the packaging ATPase genes. (separate excel file)

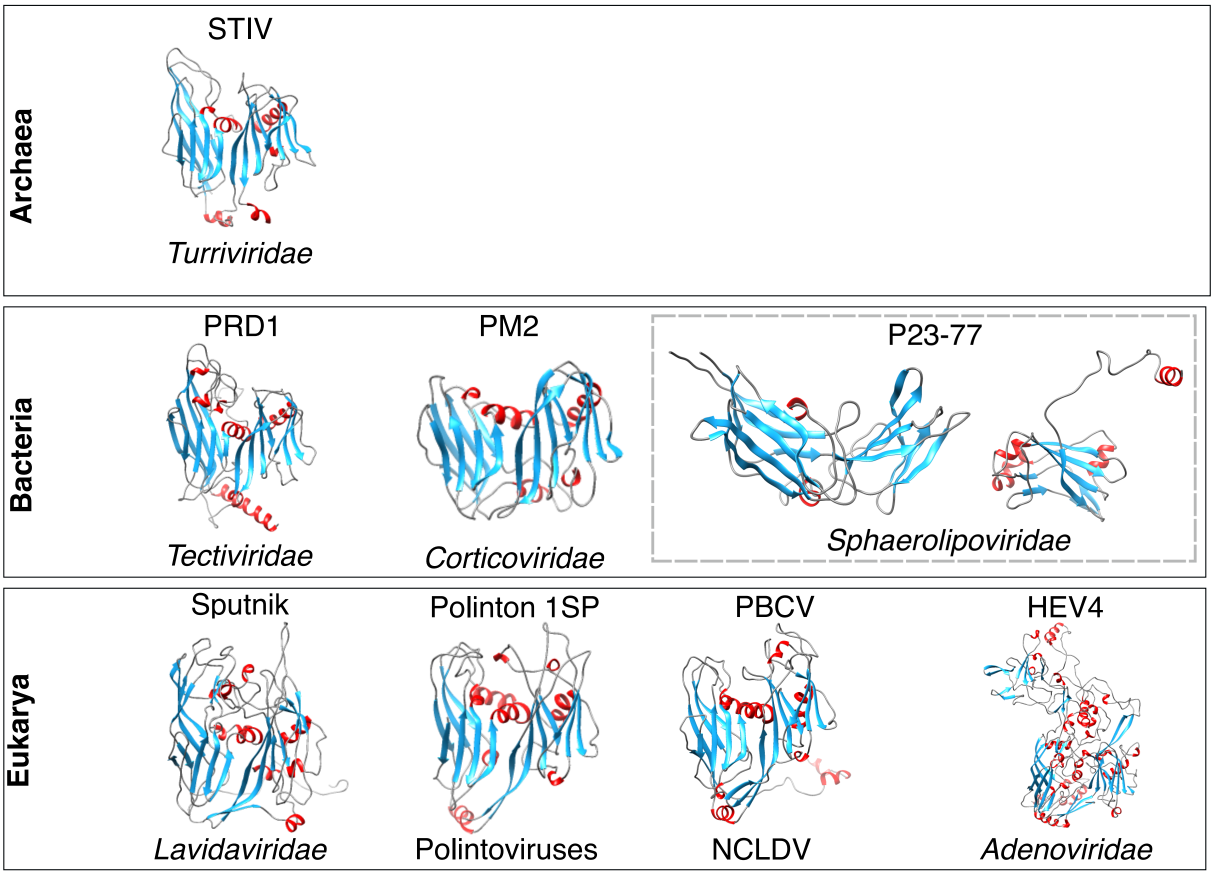


**Fig. S1. Major capsid proteins of the PRD1-adenovirus lineage.** Major capsid proteins (MCPs) with the double jelly-roll (DJR) fold and single jelly-roll (SJR) fold (boxed in dashed lines) are arranged horizontally according to the domain of life to which their hosts are classified. Virus names are provided above the structural models of various families of bacterial, archaeal and eukaryotic viruses. The structures are coloured according to the secondary structure elements: α-helices in red; β-strands in blue; and random coil in grey. The X-ray structure of the major capsid protein of Polinton 1 SP is not available and is represented with homology-based model. Sputnik, Sputnik virophage; Polinton 1 SP, Polinton 1 Strongylocentrotus purpuratus; PBCV, Paramecium bursaria Chlorella virus; NCLDV, nucleo-cytoplasmic large DNA viruses; HAE4, Human adenovirus E4; PRD1, Enterobacteria phage PRD1; PM2, Pseudoalteromonas phage PM2; STIV, Sulfolobus turreted icosahedral virus. P23-77, Thermus virus P23-77. RCSB Protein Data Bank (PDB) accession numbers for the major capsid protein structures: Sputnik, PDB entry 3J26; PBCV, PDB entry 5TIP; HAE4, PDB entry 2BVI; PRD1, PDB entry 1HB5; PM2, PDB entry 2W0C; STIV, PDB entry 2BBD; P23-77, PDB entry 3ZMN (left) and 3ZN4 (right).

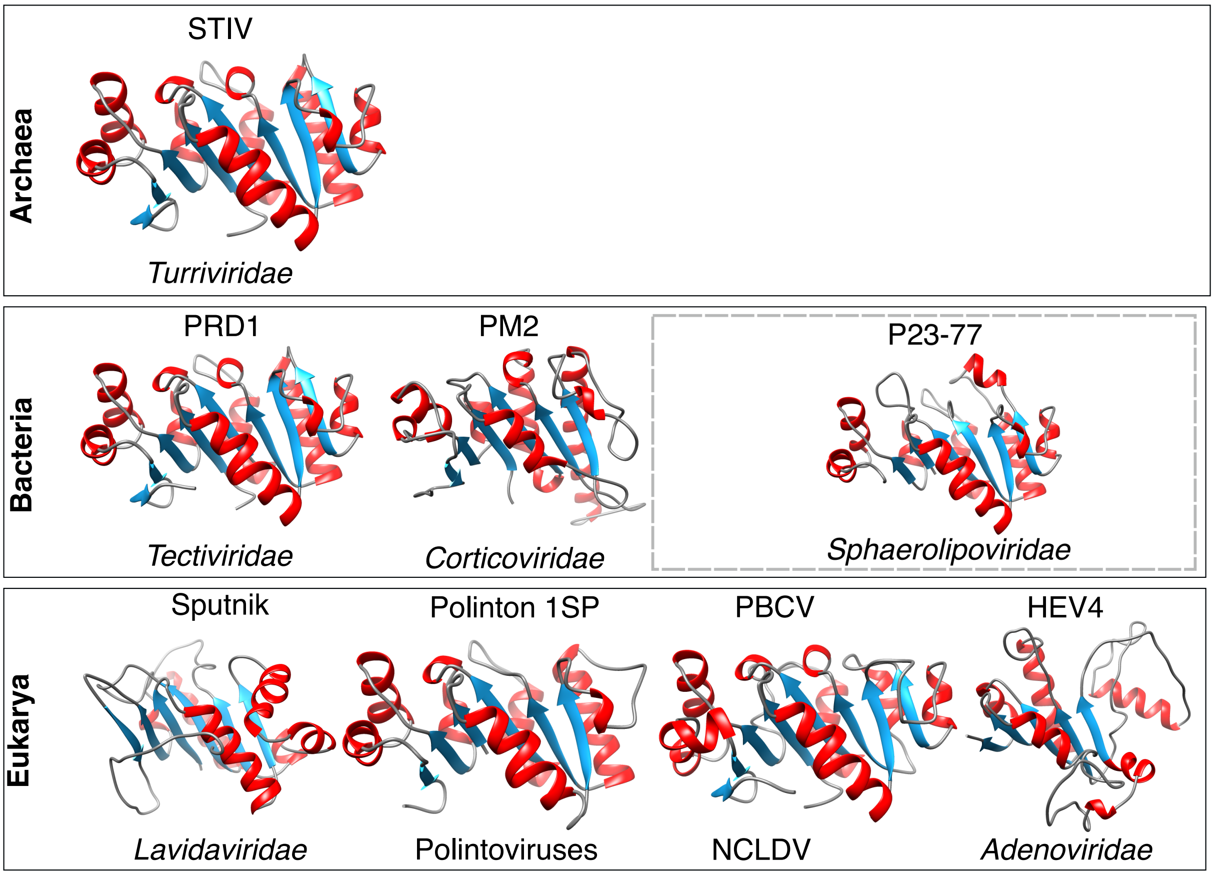


**Fig. S2. Packaging ATPases of the PRD1-adenovirus lineage.** Packaging ATPases (pATPases) are arranged horizontally according to the domain of life to which their hosts are classified. Virus names are provided above the structural models of various families of bacterial, archaeal and eukaryotic viruses. The structures are coloured according to the secondary structure elements: α-helices in red; β-strands in blue; and random coil in grey. The X-ray structures of the pATPase proteins of Sputnik, Polinton 1 SP, PBCV, HAE4, PRD1, PM2 and P23-77 are not available and are represented with homology-based models. Sputnik, Sputnik virophage; Polinton 1 SP, Polinton 1 Strongylocentrotus purpuratus; PBCV, Paramecium bursaria Chlorella virus; NCLDV, nucleo-cytoplasmic large DNA viruses; HAE4, Human adenovirus E4; PRD1, Enterobacteria phage PRD1; PM2, Pseudoalteromonas phage PM2; STIV, Sulfolobus turreted icosahedral virus. P23-77, Thermus virus P23-77. RCSB Protein Data Bank (PDB) accession numbers for the pATPase structure: STIV, PDB entry 4KFU.

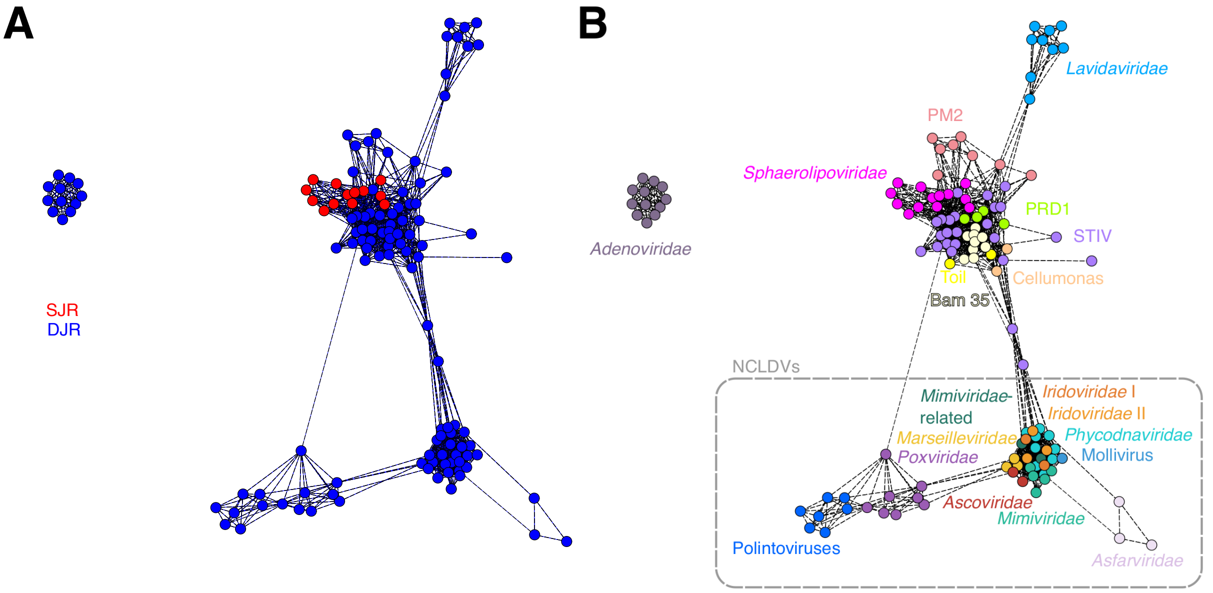


**Fig. S3. Sequence similarity networks of the PRD1-adenovirus packaging ATPase proteins.** Protein sequences were clustered using SiLiX based on the sequence similarity. Different clusters of viruses are shown as clouds of connected colored circles, with each circle corresponding to a single pATPase protein. Members are classified according to (**A**) the structure of the MCP or (**B**) the different families/subfamilies which they belong to. Abbreviations: DJR, double jelly-roll; SJR: single jelly-roll; NCLDV, Nucleo-Cytoplasmic Large DNA Viruses; STIV, Sulfolobus turreted icosahedral virus

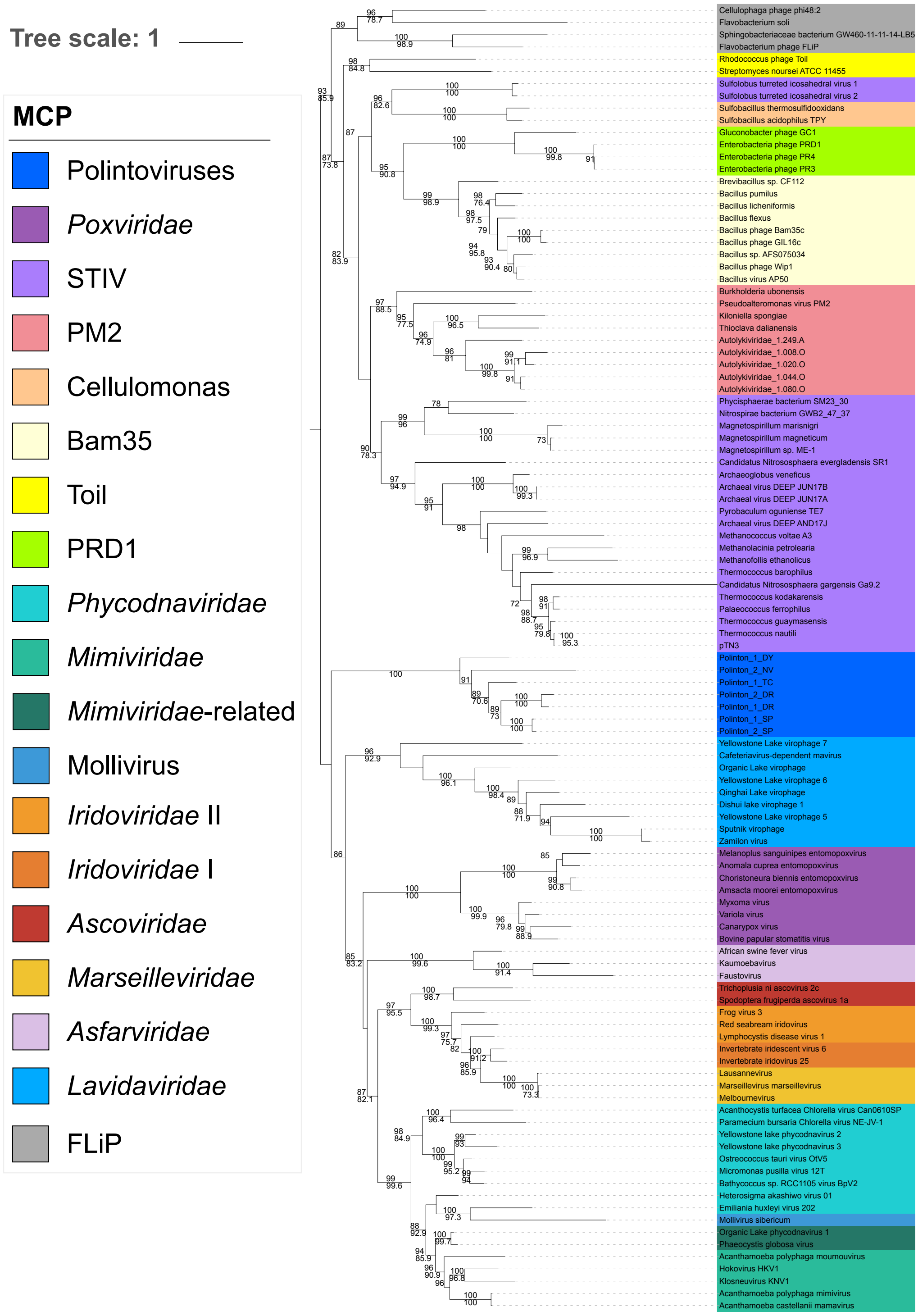


**Fig. S4. Maximum likelihood (ML) phylogenetic tree of the major capsid protein gene of the viruses from the PRD1-adenovirus lineage.** The root of the ML phylogenetic tree was between the prokaryotic and eukaryotic members. The scale-bar indicates the average number of substitutions per site. Values on top and below branches represent support calculated by ultrafast bootstrap approximation (UFBoot; 1,000 replicates) and SH-like approximate likelihood ratio test (aLRT; 1,000 replicates), respectively. Only values superior to 70 are shown. The best-fit model was LG + F + R4, which was chosen according to Bayesian Information Criterion (BIC). The alignment has 107 sequences with 262 positions.

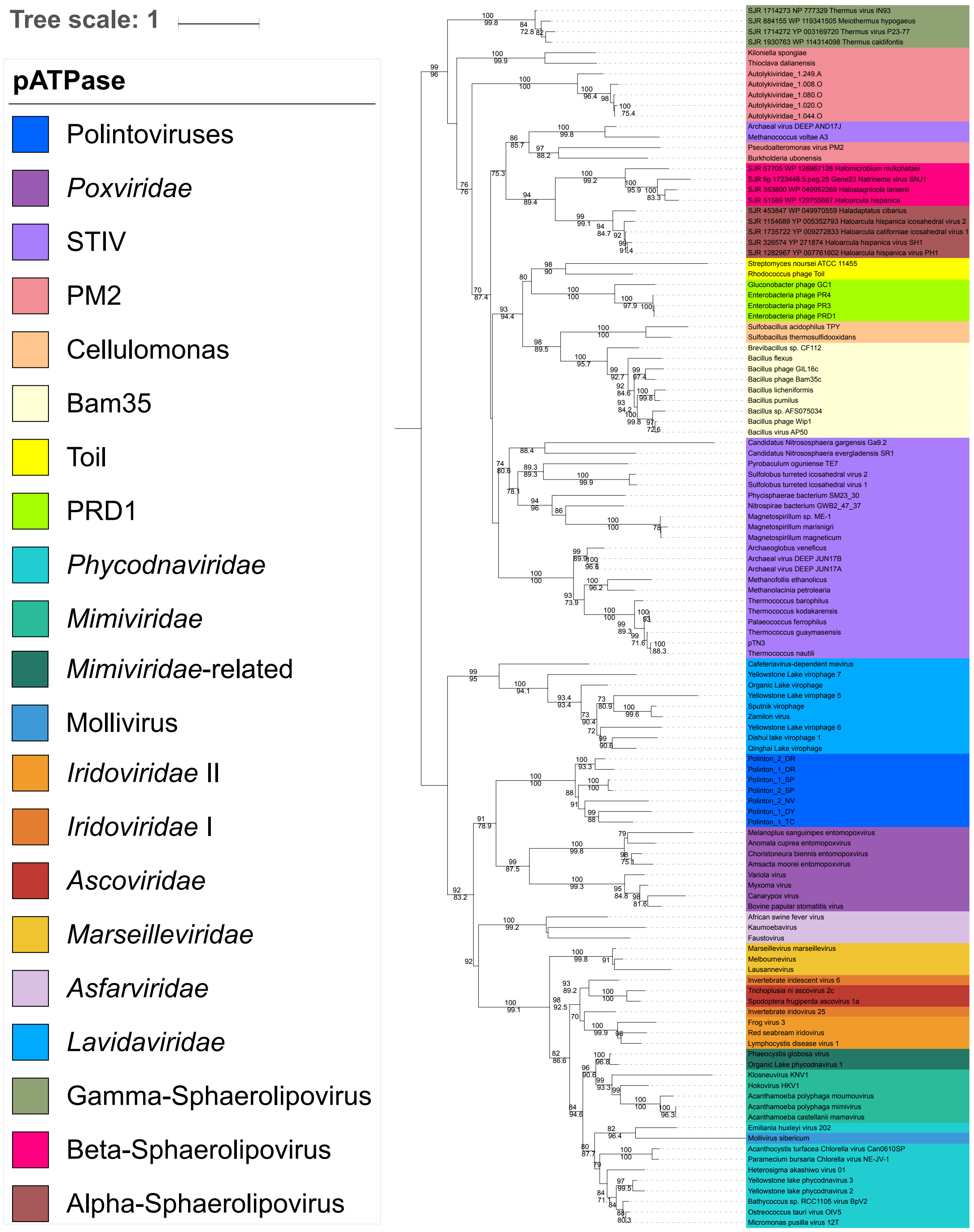


**Fig. S5. Maximum likelihood (ML) phylogenetic tree of the packaging ATPase gene of the viruses from the PRD1-adenovirus lineage.** The root of the ML phylogenetic tree was between the prokaryotic and eukaryotic members. The scale-bar indicates the average number of substitutions per site. Values on top and below branches represent support calculated by ultrafast bootstrap approximation (UFBoot; 1,000 replicates) and SH-like approximate likelihood ratio test (aLRT; 1,000 replicates), respectively. Only values superior to 70 are shown. The best-fit model was LG + R6, which was chosen according to Bayesian Information Criterion (BIC). The alignment has 116 sequences with 151 positions.

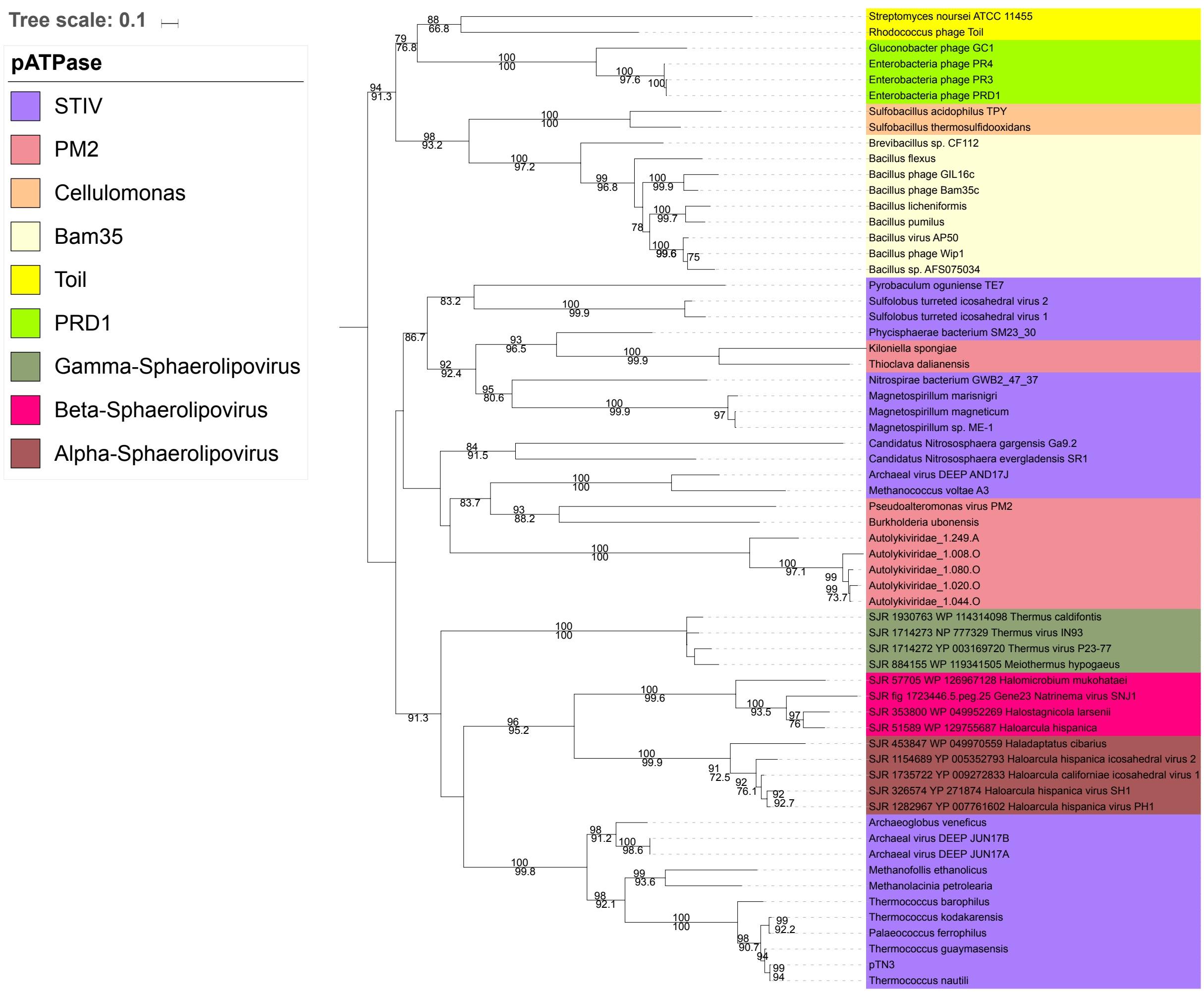


**Fig. S6. Maximum likelihood (ML) phylogenetic tree of the packaging ATPase gene of the archaeoviruses and bacterioviruses from the PRD1-adenovirus lineage.** The ML phylogenetic tree was rooted with the members of the *Tectiviridae*, which includes the PRD1, the Bam35 and the Toil. The scale-bar indicates the average number of substitutions per site. Values on top and below branches represent support calculated by ultrafast bootstrap approximation (UFBoot; 1,000 replicates) and SH-like approximate likelihood ratio test (aLRT; 1,000 replicates), respectively. Only values superior to 70 are shown. The best-fit model was LG + R4, which was chosen according to Bayesian Information Criterion (BIC). The alignment has 62 sequences with 185 positions.

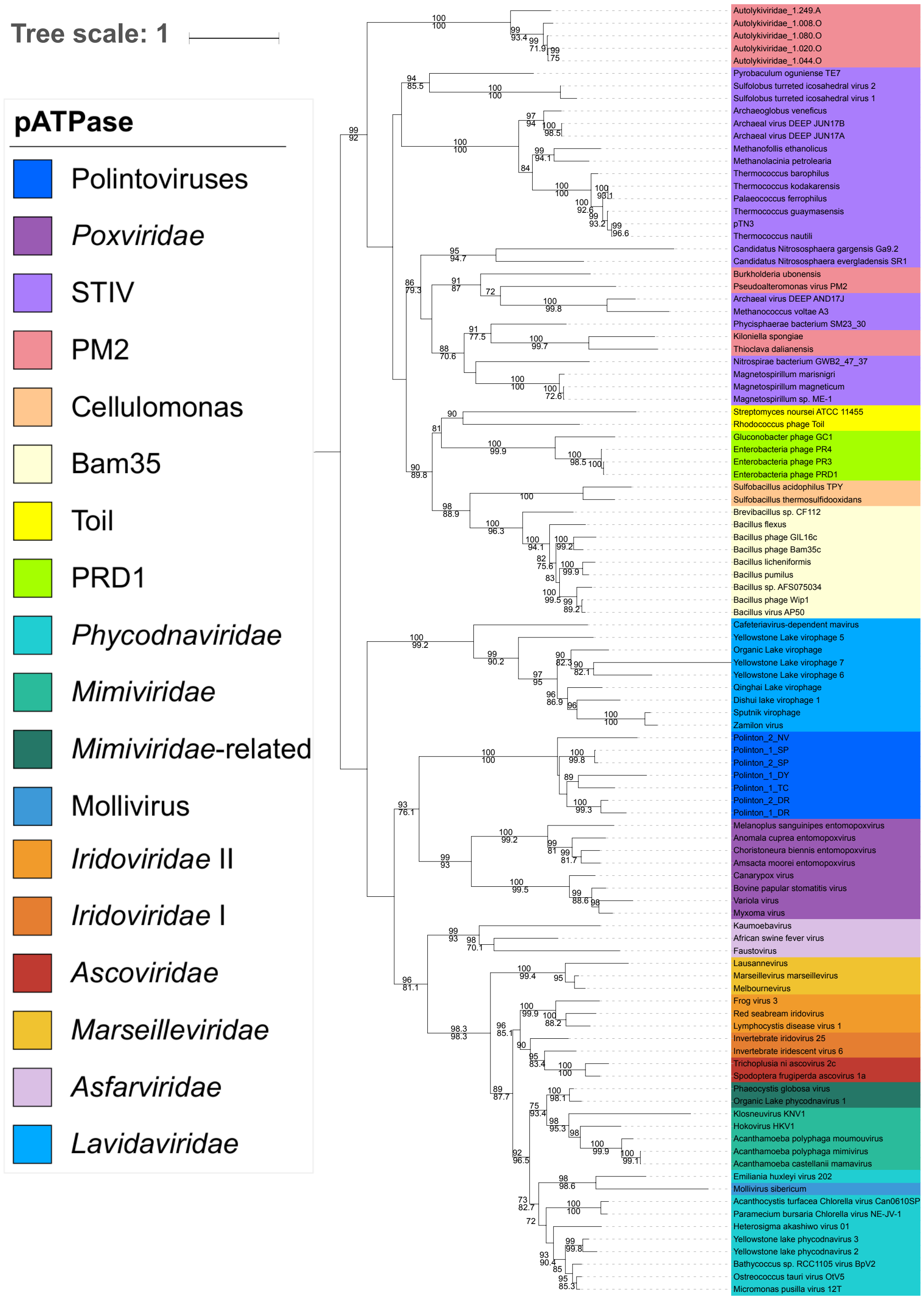


**Fig. S7. Maximum likelihood (ML) phylogenetic tree of the packaging ATPase gene of the double jelly-roll viruses from the PRD1-adenovirus lineage, excluding single jelly-roll viruses.** The root of the ML phylogenetic tree was between the prokaryotic and eukaryotic members. The scale-bar indicates the average number of substitutions per site. Values on top and below branches represent support calculated by ultrafast bootstrap approximation (UFBoot; 1,000 replicates) and SH-like approximate likelihood ratio test (aLRT; 1,000 replicates), respectively. Only values superior to 70 are shown.  The best-fit model was LG + F+ R6, which was chosen according to Bayesian Information Criterion (BIC). The alignment has 103 sequences with 171 positions.

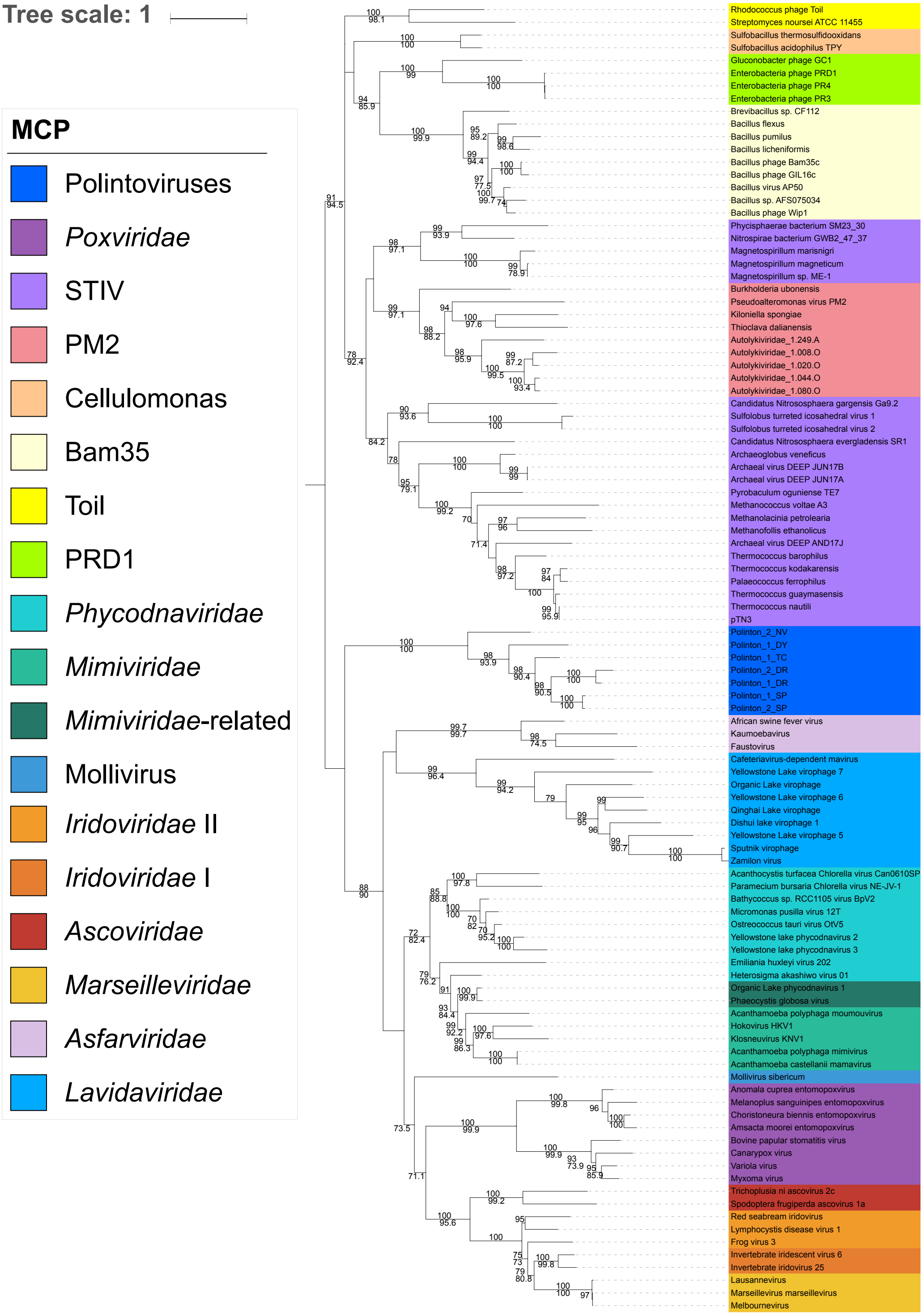


**Fig. S8. Maximum likelihood (ML) phylogenetic tree of the major capsid protein gene of the double jelly-roll viruses from the PRD1-adenovirus lineage, excluding the FLiP members.** The root of the ML phylogenetic tree was between the prokaryotic and eukaryotic members. The scale-bar indicates the average number of substitutions per site. Values on top and below branches represent support calculated by ultrafast bootstrap approximation (UFBoot; 1,000 replicates) and SH-like approximate likelihood ratio test (aLRT; 1,000 replicates), respectively. Only values superior to 70 are shown. The best-fit model was LG + F + R4, which was chosen according to Bayesian Information Criterion (BIC). The alignment has 103 sequences with 237 positions.

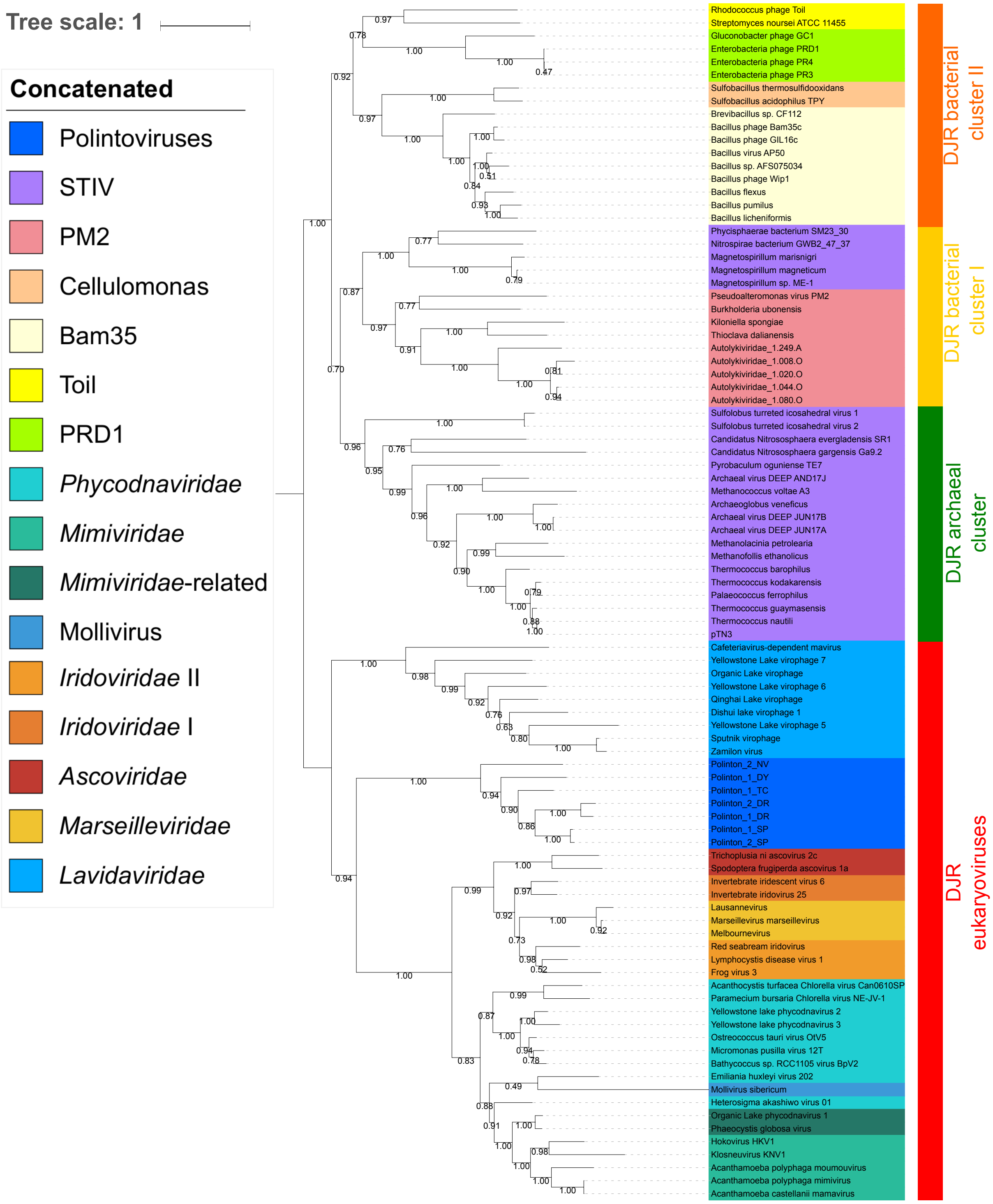


**Fig. S9. Maximum likelihood (ML) phylogenetic tree of the concatenated major capsid protein and packaging ATPase genes, excluding the *Poxviridae* and the *Asfarviridae*.** The ML phylogenetic tree was rooted between the prokaryotic and eukaryotic members. The scale-bar indicates the average number of substitutions per site. Values on the branches represent transfer bootstrap expectation (TBE) support calculated by nonparametric bootstrap (1,000 replicates). The best-fit model was LG + F + R4, which was chosen according to Bayesian Information Criterion (BIC). The alignment has 92 sequences with 425 positions.

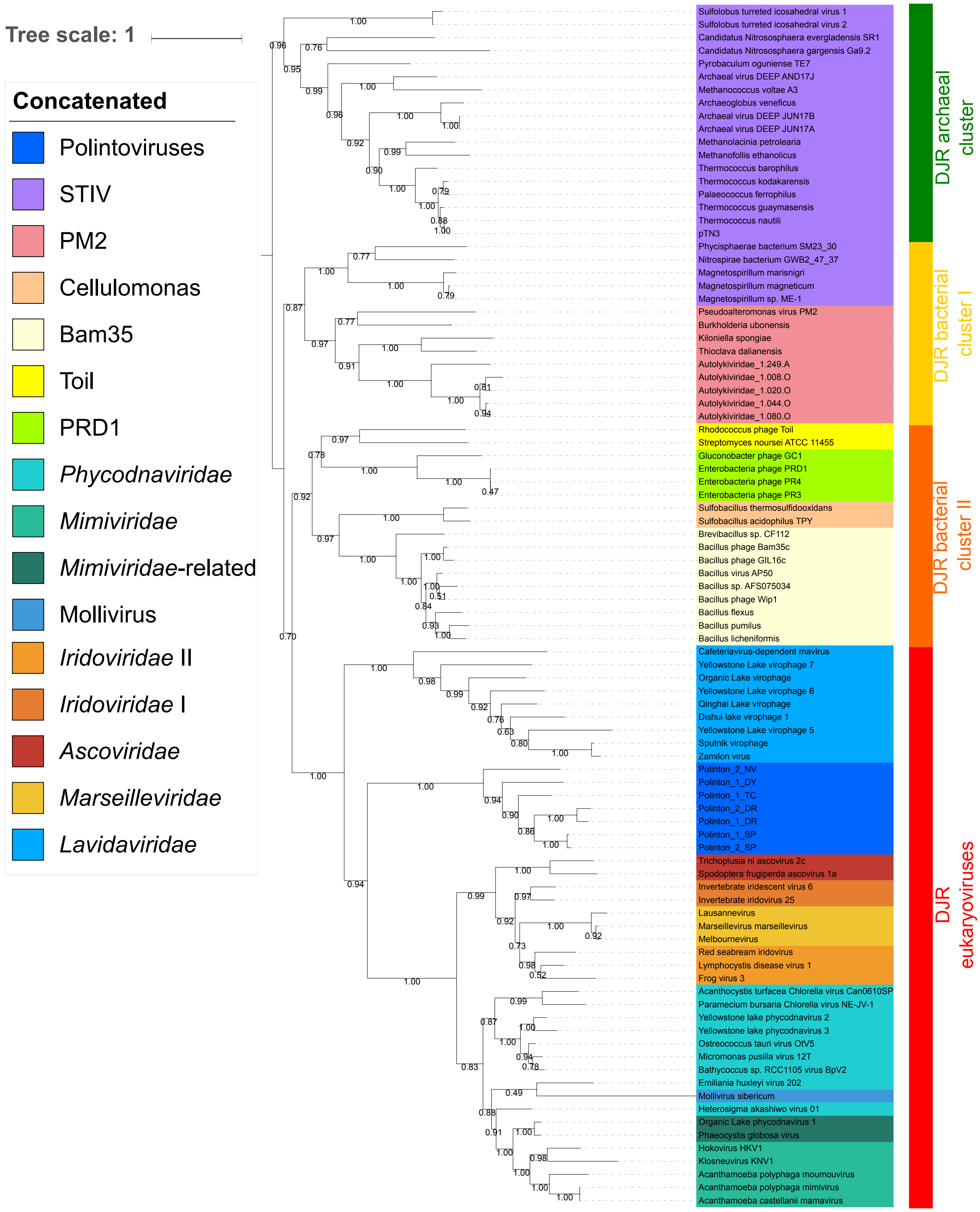


**Fig. S10. Maximum likelihood phylogenetic tree of the concatenated major capsid protein and packaging ATPase genes, rooting with the DJR archaeal cluster.** The scale-bar indicates the average number of substitutions per site. Values on the branches represent transfer bootstrap expectation (TBE) support calculated by nonparametric bootstrap (1,000 replicates). The best-fit model was LG + F + R4, which was chosen according to Bayesian Information Criterion (BIC). The alignment has 92 sequences with 425 positions.

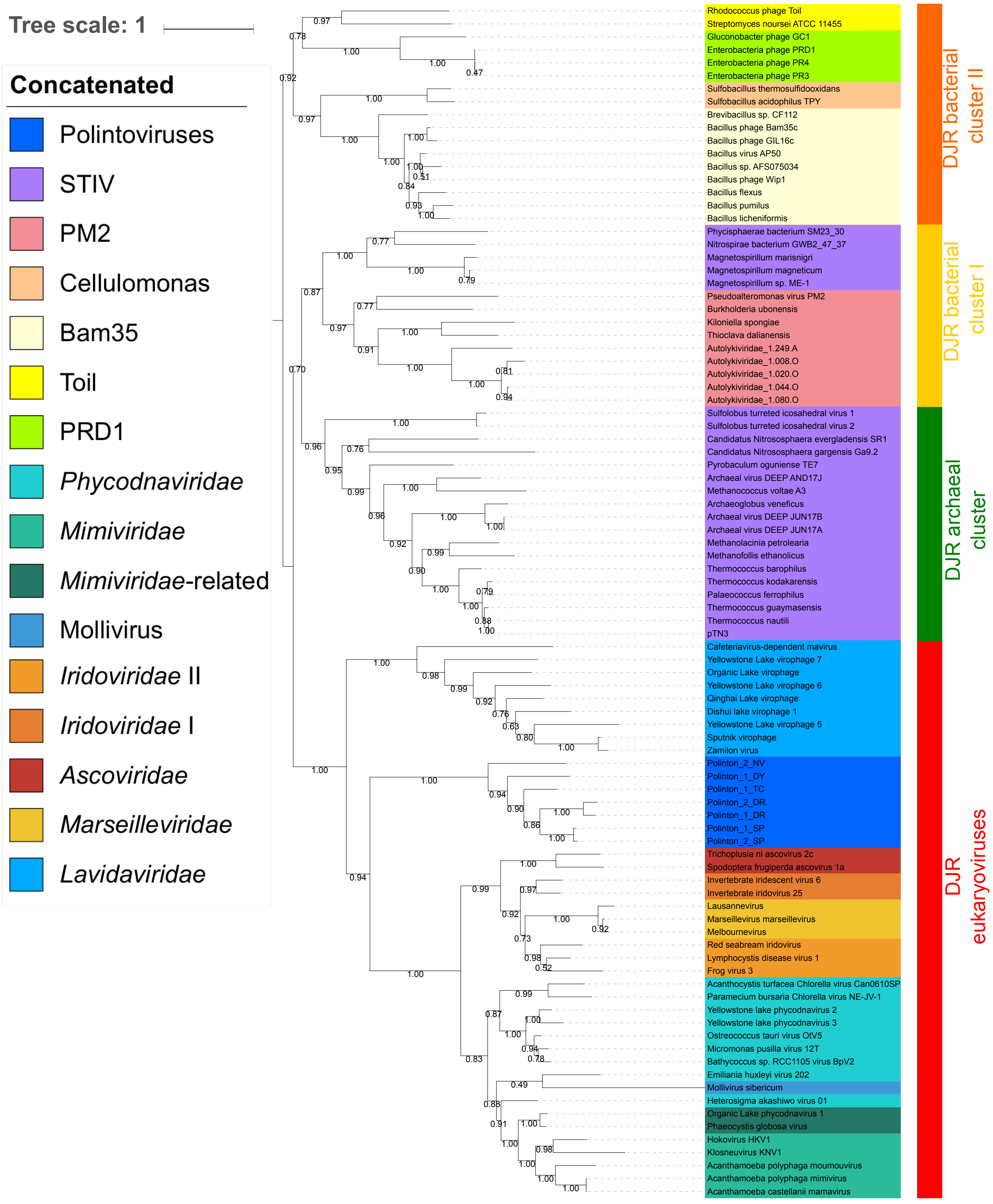


**Fig. S11. Maximum likelihood phylogenetic tree of the concatenated major capsid protein and packaging ATPase genes, rooting with the DJR bacterial cluster II.** The scale-bar indicates the average number of substitutions per site. Values on the branches represent transfer bootstrap expectation (TBE) support calculated by nonparametric bootstrap (1,000 replicates). The best-fit model was LG + F + R4, which was chosen according to Bayesian Information Criterion (BIC). The alignment has 92 sequences with 425 positions.

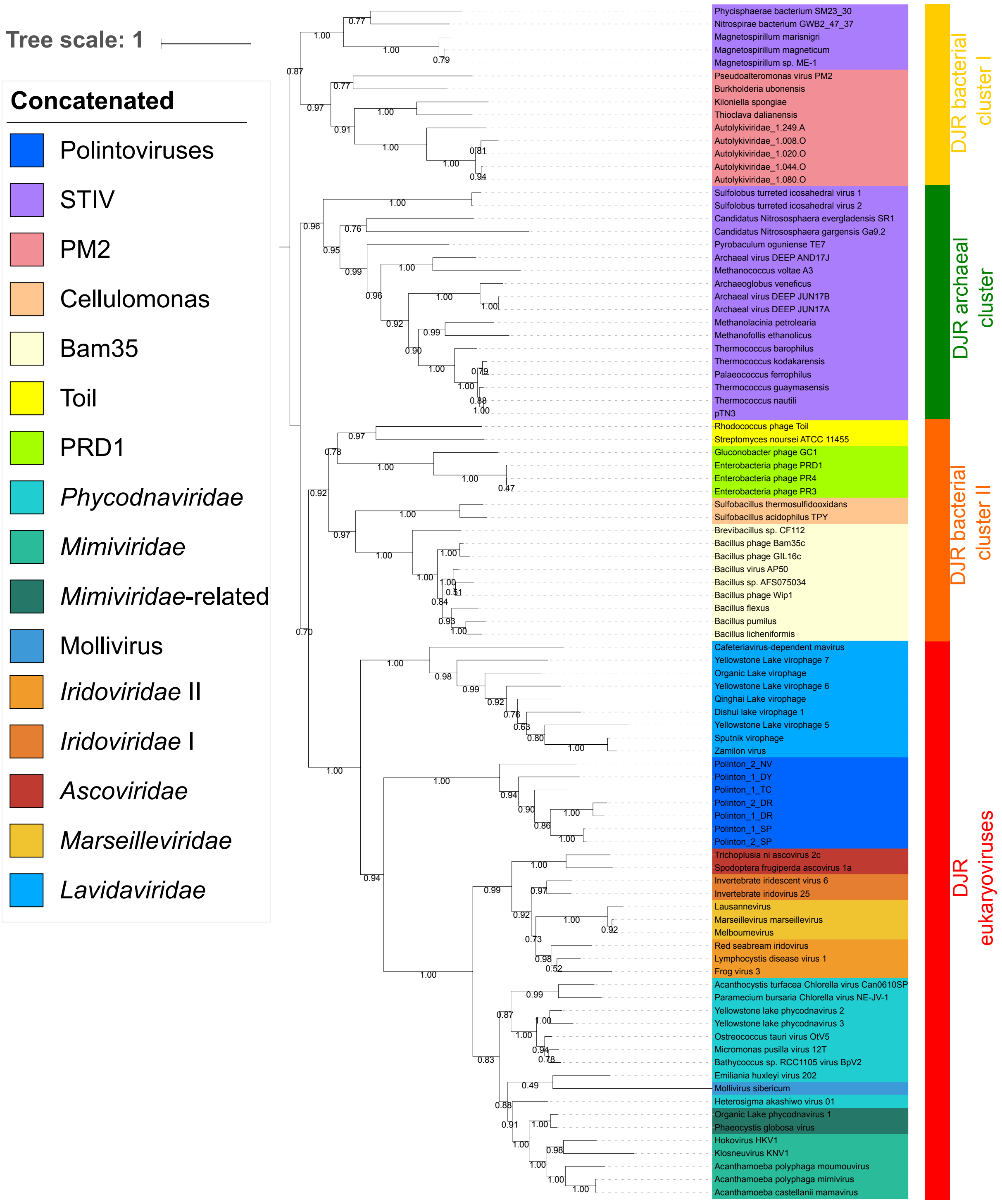


**Fig. S12. Maximum likelihood phylogenetic tree of the concatenated major capsid protein and packaging ATPase genes, rooting with the DJR cluster I.** The scale-bar indicates the average number of substitutions per site. Values on the branches represent transfer bootstrap expectation (TBE) support calculated by nonparametric bootstrap (1,000 replicates). The best-fit model was LG + F + R4, which was chosen according to Bayesian Information Criterion (BIC). The alignment has 92 sequences with 425 positions.

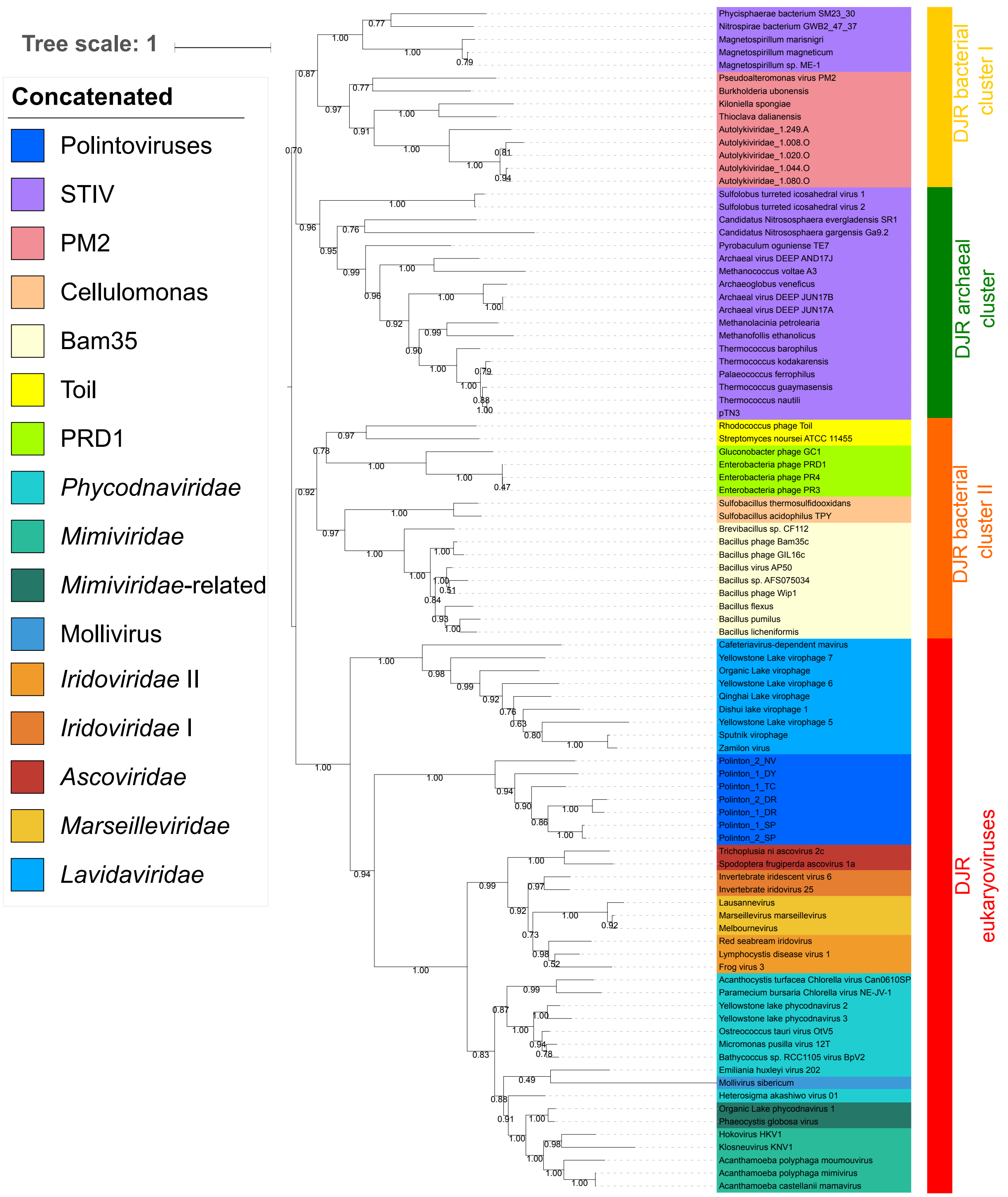


**Fig. S13. Maximum likelihood phylogenetic tree of the concatenated major capsid protein and packaging ATPase genes, rooting between the DJR archaeal cluster and the DJR bacterial cluster II.** The scale-bar indicates the average number of substitutions per site. Values on the branches represent transfer bootstrap expectation (TBE) support calculated by nonparametric bootstrap (1,000 replicates). The best-fit model was LG + F + R4, which was chosen according to Bayesian Information Criterion (BIC). The alignment has 92 sequences with 425 positions.
